# Mitochondrial Respiratory Chain Function is crucial for Muscle Toxicity in Facioscapulohumeral Muscular Dystrophy

**DOI:** 10.1101/2025.11.25.690559

**Authors:** Philipp Heher, Jakob Pfeiffer-Vogl, Massimo Ganassi, Raul-Cristian Fulea, Liam McGuire, Elise N Engquist, Magda Tyszkiewicz, Johanna Prueller, Johannes Grillari, Giorgio Tasca, Christopher RS Banerji, Peter S Zammit

**Affiliations:** Randall Centre for Cell and Molecular Biophysics, King’s College London, Guy’s Campus, London, UK; John Walton Muscular Dystrophy Research Centre, NIHR Newcastle Biomedical Research Centre, Newcastle University and Newcastle Hospitals NHS Foundation Trust, Newcastle upon Tyne, UK; Department of Clinical Pharmacology, Medical University of Vienna, Vienna, Austria; Genetic Therapy Accelerator Centre, Queen Square Institute of Neurology, University College London, London, UK; Cardiovascular and Genomics Research Institute, City St. George’s, University of London, London, UK; Ludwig Boltzmann Institute for Traumatology. The Research Center in Cooperation with AUVA, Vienna, Austria; Austrian Cluster for Tissue Regeneration, Vienna, Austria; Institute of Molecular Biotechnology, BOKU University, Vienna, Austria; Comprehensive Cancer Centre, King’s College London, Guy’s Campus, London, UK; The Alan Turing Institute, London, UK; University College London NHS Trust, London, UK

**Keywords:** FSHD, facioscapulohumeral muscular dystrophy, mitochondria, mitochondrial respiratory chain, reverse electron transfer, DUX4

## Abstract

Facioscapulohumeral muscular dystrophy (FSHD) is driven by DUX4-induced toxicity, yet the pathomechanisms remain unclear. Here, we identify persistent transcriptional suppression of the mitochondrial respiratory chain and metabolic rewiring in FSHD muscle biopsies and myotubes. Using DUX4-inducible human myogenic cells, we show that DUX4 target gene activation is accompanied by mitochondrial function impairment and Caspase 9-mediated apoptosis. Reverse electron transfer (RET) at complex I is the dominant oxidative stress generating mechanism in FSHD muscle cells. RET-driven mitochondrial reactive oxygen species (mitoROS) are the trigger for oxidative stress and apoptosis, which is a unique feature of FSHD mitochondria. Pharmacological inhibition of RET suppresses mitoROS, reduces Caspase 9 activation, and rescues abnormal myogenesis in FSHD cell lines. Importantly, DUX4 is non-toxic in oxidative phosphorylation-deficient human myogenic cells. Our findings identify the mitochondrial respiratory chain as a key mediator of DUX4 toxicity and highlight RET inhibition as a potential therapeutic approach for FSHD.

**Graphical abstract:** (Created with BioRender.com)

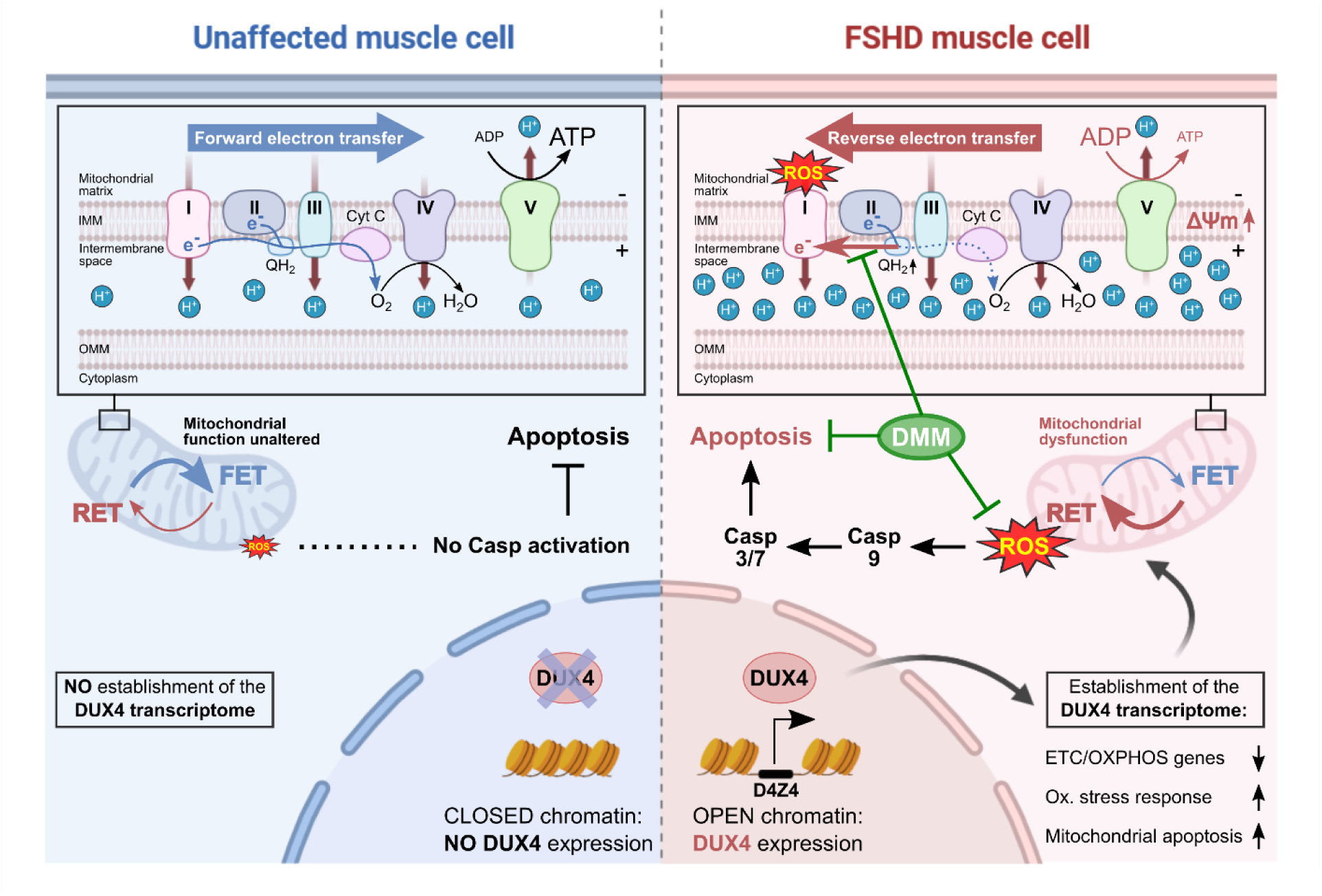

## INTRODUCTION

Skeletal muscle has high but variable energy demands and thus needs to be able to rapidly adapt to metabolic challenges such as those posed by cycles of exercise/inactivity and nutrient availability. Muscle contraction, where consumption of adenosine triphosphate (ATP) can increase up to 100-fold [1], necessitates rapid energy provision. Mitochondria are critical players in these dynamics, providing most of the energy required for muscle function through the concerted actions of the electron transport chain (ETC), oxidative phosphorylation (OXPHOS), the tricarboxylic acid (TCA) cycle and fatty acid β-oxidation. Mitochondria are also major cell signalling hubs controlling pathways including redox signalling through mitochondrial reactive oxygen species (mitoROS), the intrinsic apoptosis pathway, mitochondrial Ca^2+^ signalling, adaptation to hypoxia, as well as many other anterograde (nucleus-to-mitochondria) and retrograde (mitochondria-to-nucleus) signal transduction cascades [2]. Contribution to these pathways establishes a central role of mitochondrial quality control in skeletal muscle homeostasis. Tightly coordinated interplay between mitochondrial dynamics (i.e. balance between mitochondrial fusion and fission), biogenesis, proteostasis and mitophagy maintains mitochondrial health and contributes to efficient skeletal muscle function [3]. Signalling cascades governing these pathways are a complex biochemical integration that requires participation of multiple cellular components including mitochondria, nucleus, cell membrane, ribosome, lysosome and proteasome, perturbation of this cellular bioenergetic system has broad consequences at virtually all levels of skeletal muscle homeostasis [4]. This complexity is demonstrated in mitochondrial disorders, where mutations in mitochondrial proteins encoded in the nuclear or mitochondrial genome usually affect numerous organs [5], particularly skeletal muscle in the form of mitochondrial myopathy [6].

This carefully coordinated system of metabolic adaptation is also perturbed in several muscle diseases. Muscular Dystrophies are a heterogenous group of genetic disorders leading to progressive muscle weakness and wasting [7]. Mitochondrial dysfunction and oxidative stress are common hallmarks of most, if not all, muscular dystrophies including Duchenne and Becker muscular dystrophy (DMD and BMD, respectively), myotonic dystrophy type 1, limb girdle muscular dystrophies and facioscapulohumeral muscular dystrophy (FSHD) [8]. However, it is unclear whether metabolic stress is a causal pathomechanistic feature or predominantly a non-specific consequence of different damaging processes involved in muscle damage and degeneration, as mechanisms linking altered mitochondrial metabolism with aetiology of oxidative stress and damage remain poorly understood.

FSHD is the third most common muscular dystrophy [9]. It presents with slowly progressing, asymmetric muscle weakness and atrophy, usually starting in facial, shoulder girdle and upper limb muscles, before progressing to lower limb and core muscles [10].

Aberrant expression of the pioneer transcription factor *Double Homeobox 4 (DUX4)* in skeletal muscle is the driver of pathology [11]. This mis-expression originates from a macrosatellite array consisting of *D4Z4* units in the subtelomere of chromosome 4 (4q35), with each D4Z4 repeat harbouring an open reading frame encoding the *DUX4* retrogene. Epigenetic de-repression of the D4Z4 array, in combination with a polyadenylation signal (PAS) on disease-permissive 4qA haplotypes, leads to stable expression of *DUX4* from the distal-most *D4Z4* unit [12, 13]. In ∼95% of FSHD cases (FSHD1; OMIM: 158900), epigenetic derepression is mediated through contraction of the *D4Z4* repeat array, from >100-11 *D4Z4* units typical of unaffected individuals in the Western world, to 10-1 copies on at least one allele in FSHD1 [14–16]. In the residual ∼5% (FSHD2; OMIM: 158901), epigenetic de-repression of usually ∼20-11 D4Z4 units on a disease-permissive 4qA haplotype is caused by pathogenic variants in genes involved in methylation/epigenetic regulation of the region [17–20].

*DUX4* misexpression in skeletal muscle is linked to oxidative stress *in vitro* and *in vivo*. After the seminal observation that FSHD patient-derived muscle cells have an usually high susceptibility to apoptotic cell death under exogenous oxidative stress [21], perturbations of cellular bioenergetics and the oxidative stress response have been identified [22–26]. Other studies showed that FSHD patients exhibit mitochondrial dysfunction and redox imbalances with possible correlations between oxidative damage and muscle functional impairment [27]. These findings suggest that mitochondrial dysfunction is a driver of oxidative stress in FSHD. Transcriptomic meta-analysis and proteomic studies confirmed dysregulation of mitochondrial function and oxidative metabolism in *in vitro* FSHD models and patient biopsies [28], and that the FSHD proteome is characterised by altered mitochondrial turnover and metabolic reprogramming [29, 30]. In addition, we recently showed that repression of mitochondrial biogenesis through Peroxisome Proliferator-Activated Receptor Gamma, Coactivator 1 Alpha (PGC1α) contributes to impaired myogenesis in FSHD [31]. We also found that DUX4 causes mitochondrial complex I dysfunction, alongside mitochondrial membrane potential (ΔΨm)-driven enhanced mitoROS production [32]. Furthermore, transcriptional repression of protein coding genes in the mitochondrial genome characterises FSHD muscle biopsies [32, 33]. DUX4 itself exerts direct toxicity in muscle through oxidative stress [34], for example through induction of oxidative DNA damage [35]. Interestingly, while very low levels of DUX4 trigger enhanced mitoROS production in muscle cells [36], and in FSHD patient iPSC-derived muscle cells, *DUX4* itself is subject to ROS-mediated redox regulation [37]. Thus, metabolic stress is an integral component of FSHD pathology and points to direct mitochondrial involvement in DUX4-mediated oxidative stress and muscle cell apoptosis.

Here, we show that the mitochondrial respiratory chain is central to DUX4-induced oxidative stress. We identify transcriptional impairment of the mitochondrial respiratory chain and mitochondrial apoptosis in FSHD muscle biopsies. We pinpoint OXPHOS defects in FSHD myotubes that consistently show an altered balance between glycolytic and mitochondrial ATP generation. Using doxycycline (DOX)-inducible *DUX4* human myoblasts (iDUX4), we demonstrate that DUX4 target gene activation marks onset of mitochondrial functional impairment and simultaneous Caspase 9 (Casp9) activation. Furthermore, we show that the mitochondrial respiratory chain is the major ROS producing system in DUX4-expressing and FSHD patient-derived muscle cells, and that reverse electron transfer (RET) is the mitoROS generating mechanism upstream of Casp9 activation. Importantly, RET-dependent ROS production is specific to FSHD mitochondria. Suppression of RET at complex II reduces oxidative stress through mitoROS, abolishes Casp9-mediated apoptosis and rescues abnormal FSHD myogenesis. We confirm this mechanism by demonstrating that DUX4 is non-toxic in OXPHOS-deficient iDUX4 muscle cells. Summarising, RET in the mitochondrial respiratory chain is central to relaying DUX4-induced toxicity, and our work clarifies the oxidative stress mechanism in FSHD muscle cells that is druggable with compounds acting directly on the respiratory chain.

## RESULTS

### FSHD myotubes are characterised by a global ATP deficit and altered balance between glycolytic and OXPHOS ATP production

DUX4 is known to impair OXPHOS, resulting in significantly higher mitoROS levels in DUX4-overexpressing and FSHD-derived muscle cells [32]. We initially assessed mitochondrial function using respirometry in three paired FSHD cellular models derived from patients with either somatic mosaicism for the contracted D4Z4 allele (where lines ‘54-12’ and ‘K8’ have a contracted allele and ‘54-6’ and ‘K4’ are their D4Z4 uncontracted isogenic controls) or from patients with a contracted D4Z4 allele (‘16A’) and a sibling-matched control (‘16U’; Fig. 1A).

**Fig. 1:**
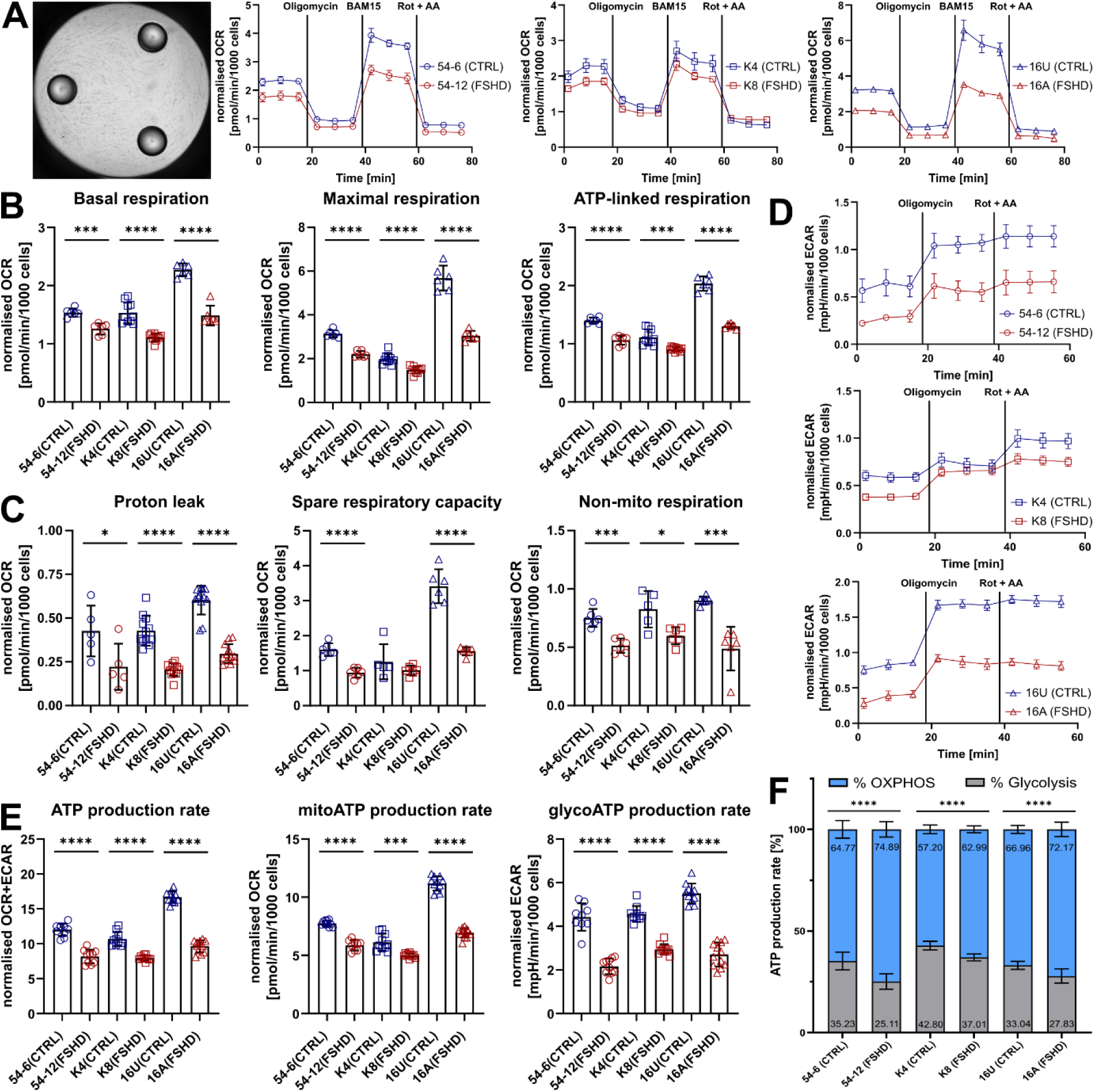
FSHD myotubes are characterised by an ATP deficit and altered metabolic setup. **(A)** Left: Brightfield micrograph of an FSHD patient-derived myotube culture after Seahorse respirometry. Right: normalised oxygen consumption rate (OCR) curves of myotubes derived from three independent FSHD patient cell models (54-12, K8, 16A), alongside matched unaffected controls (54-6, K4, 16U). **(B)** Impairment of mitochondrial basal, maximal and ATP-linked respiratory function in FSHD myotubes. **(C)** Altered proton leak, spare respiratory capacity and non-mitochondrial (“Non-mito”) respiration in FSHD myotubes. **(D)** Normalised extracellular acidification rate (ECAR) curves of the three FSHD myotube models, alongside unaffected controls. **(E)** ATP production rates demonstrating an ATP deficit in FSHD myotubes arising from both defective mitochondrial and glycolytic ATP generating systems (mitoATP and glycoATP, respectively), resulting in **(F)** an altered balance between mito and glycoATP production. [n=6-10, data is mean ± s.d., where \**p*<0.05, \*\*\**p*<0.001, \*\*\*\**p*<0.0001].

FSHD myotubes (54-12, K8 and 16A) showed significantly reduced basal, maximal and ATP-linked respiration rates compared to isogenic or sibling-matched controls (54-6, K4, 16U; Fig. 1B). Proton leaks, spare respiratory capacity and non-mitochondrial respiration were also altered (Fig. 1C), suggesting widespread mitochondrial functional changes in FSHD myotubes. To determine how these changes affected ATP production, we also measured ATP production rates, with a focus on relative glycolytic versus OXPHOS ATP rates (Fig. 1D). As expected, total ATP production rates where significantly lower in FSHD myotubes, but both reduced mitochondrial and glycolytic ATP production contributed to this ATP deficit (Fig. 1E). Relative contribution of OXPHOS (mitoATP) versus glycolytic ATP (glycoATP) to total ATP production was consistently higher in FSHD myotubes (Fig. 1F), suggesting that glycolytic perturbations also majorly contribute to metabolic stress in these cells.

Myotubes predominantly use OXPHOS to produce ATP, so we also assayed these parameters in myoblasts, which mostly rely on glycolytic ATP production. Interestingly, FSHD myoblasts had significantly higher mitochondrial respiration rates (Fig. S1A, B). Proton leaks, spare respiratory capacity and non-mitochondrial respiration were also consistently higher compared to controls (Fig. S1C). The same significant reduction in total ATP production rates was found in FSHD myoblasts (Fig. S1D, E). However, this ATP deficit could be attributed to reduced glycolytic ATP production, as mitoATP production was increased in FSHD myoblasts. Relative contribution of mitoATP versus glycoATP to total ATP production was consistently higher in FSHD myoblasts (Fig. S1F), in line with the findings in FSHD myotubes (Fig. 1F).

### DUX4-induced transcriptional changes mark onset of mitochondrial dysfunction and mitoROS-induced Caspase 9-dependent activation of the intrinsic apoptotic pathway

We next tested how *DUX4* expression affects mitochondrial function in myotubes using the DOX-inducible human LHCN-M2-iDUX4 (iDUX4) muscle cell model [38]. To establish a dose-dependency relationship between DUX4 and mitochondrial functional changes, iDUX4 cells were differentiated into myotubes, and respirometry was performed after 36h of *DUX4* expression at variable induction rates to achieve low, intermediate and high DUX4 levels (62.5,125 and 250 ng/mL DOX; Fig. 2A-C). *DUX4* expression in iDUX4 myotubes significantly reduced basal, maximal and ATP-linked respiration in a dose-dependent manner (Fig. 2D), and reduced coupling efficiency, spare respiratory capacity and non-mitochondrial respiration, while proton leak was affected at lower DUX4 levels (Fig. 2E). Total ATP production was reduced upon intermediate or high DUX4 induction, as mitoATP deficits after low level induction (62.5 ng/mL DOX) could be compensated by increased glycoATP production (Fig. 2F). At intermediate (125 ng/mL DOX) and high (250 ng/mL DOX) induction, mitoATP production was the predominantly affected bioenergetic circuit, while glycoATP production was unaltered. All DUX4 induction regimes increased the relative contribution of glycoATP to total ATP, at intermediate and high induction predominantly through reduction of mitoATP, suggesting that DUX4 impacts metabolic switching between glycolytic and OXPHOS ATP production mainly by affecting OXPHOS.

**Fig. 2:**
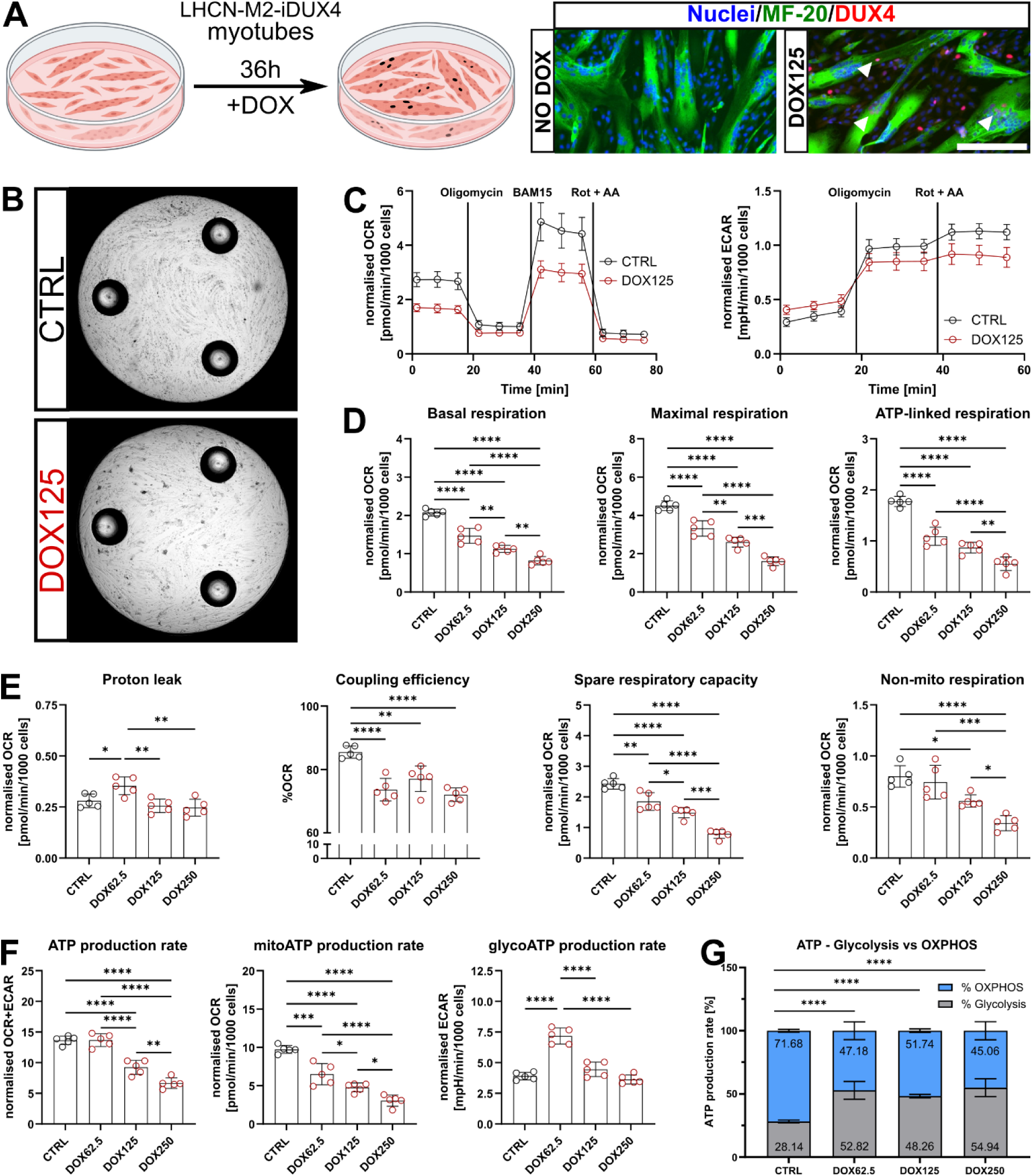
DUX4 expression in human myotubes triggers metabolic stress through mitochondrial dysfunction in a dose-dependent manner. **(A)** Left: schematic of the LHCN-M2-DUX4-inducible human myotube model (“iDUX4”), where *DUX4* expression can be induced through doxycycline (DOX). Scheme was created with BioRender.com. Right: immunofluorescence micrographs of iDUX4 myotubes expressing DUX4 after DOX (125 ng/mL) for 24h (red=DUX4, green=MF-20, blue=nuclei; scale bar=200 μm). Arrowheads mark DUX4-positive myonuclei. **(B)** Brightfield micrographs of iDUX4 control (CTRL, top) and DOX-induced (125 ng/mL DOX for 36h, bottom) myotubes after Seahorse respirometry. **(C)** Normalised oxygen consumption rate (OCR - left) and extracellular acidification rate (ECAR - right) curves of iDUX4 control (CTRL) and DOX-induced (125 ng/mL DOX for 36h) myotubes. **(D)** Dose-dependent reduction of mitochondrial basal, maximal and ATP-linked respiratory function in iDUX4 myotubes after induction of *DUX4* expression through exposure to different concentrations of DOX for 36h. **(E)** Altered proton leak, and reduced coupling efficiency, spare respiratory capacity and non-mitochondrial (“Non-mito”) respiration in iDUX4 myotubes after induction of *DUX4* through exposure to DOX for 36h. **(F)** ATP production rates demonstrating a gradual ATP deficit in iDUX4 myotubes after induction of *DUX4* through exposure to DOX for 36h, arising from defective oxidative phosphorylation (OXPHOS; mitoATP), resulting in **(F)** an altered balance between mito and glycoATP production. [n=5-6, data is mean ± s.d., where \**p*<0.05, \*\**p*<0.01, \*\*\**p*<0.001, \*\*\*\**p*<0.0001].

We also tested DUX4-induced effects on mitochondrial function in undifferentiated proliferating iDUX4 myoblasts. *DUX4* expression at variable induction rates to achieve low, intermediate and high DUX4 levels (15.62, 31.25 and 62.50 ng/mL DOX) triggers apoptosis as measured by Annexin V in a dose-dependent manner starting ∼16h after DOX administration (Fig. S2A, B), which was the timepoint chosen for respirometry analysis (Fig. S2C, D). *DUX4* expression significantly reduced mitochondrial respiration in iDUX4 myoblasts, with a dose dependent effect on ATP-linked respiration (Fig. S2E), similar to iDUX4 myotubes (Fig. 2). Proton leak and coupling efficiency correlated inversely with DOX concentration, and spare respiratory capacity was reduced at intermediate (31.25 ng/mL DOX) and high (62.50 ng/mL DOX) levels of DUX4 induction, while non-mitochondrial respiration was unaffected in myoblasts (Fig. S2F). ATP rate measurements in iDUX4 myoblasts were largely consistent with those in iDUX4 myotubes, with deficits in mitoATP production rates driving the DUX4-induced reduction in total ATP and relative increase of glycoATP (Fig. S2G, H).

Having established that DUX4 triggers mitochondrial dysfunction, rewiring metabolism towards a more glycolytic setup, we next examined the kinetics of *DUX4* expression, mitochondrial dysfunction and muscle cell apoptosis. We generated an iDUX4 reporter line carrying the promoter of the DUX4 target gene *ZSCAN4* driving tdTomato (iDUX4^ZSCAN4-tdT^) to live monitor DUX4 target gene expression (Fig S6). DUX4 protein was readily detectable via immunofluorescence 2h-post DUX4 induction (62.50 ng/mL DOX) in iDUX4^ZSCAN4-tdT^ myoblasts, with DUX4 target gene activation (tdTomato reporter fluorescence) first evident at 4h (Fig. 3A). Intriguingly, this 4h timepoint also corresponded to the onset of mitochondrial dysfunction, evident as significant reduction both in mitochondrial respiration (Fig. 3B, C) and ATP production rate (Fig. 3D, E), driven by decreased mitoATP formation in response to reduced ATP-linked respiration. Notably, ATP production through glycolysis (glycoATP) was initially not affected upon DUX4 target gene activation (Fig. 3E). At 4h post-DUX4 induction, Casp9 activation exceeded control levels, followed at 8h by downstream activation of the effector Casp3/7 (Fig. 3F). Parallel Annexin V apoptosis assaying revealed that overt muscle cell apoptosis followed later at 14h, so ∼10h-post activation of DUX4-induced mitochondrial apoptotic signalling through Casp9. Therefore, mitochondrial dysfunction and apoptotic signalling through the intrinsic pathway are very early events in response to DUX4 induction (Fig. 3G).

**Fig. 3:**
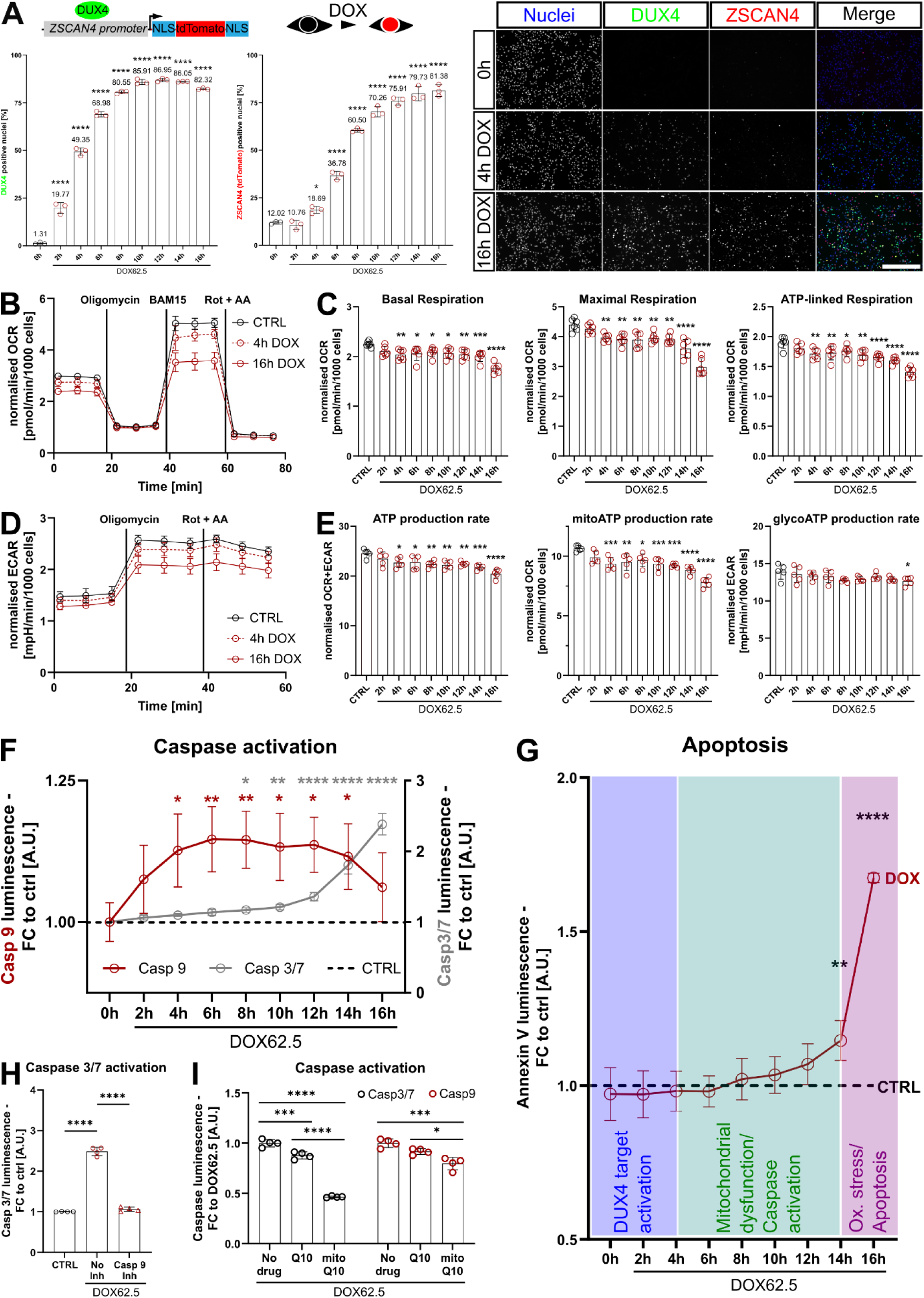
DUX4-induced apoptosis is driven by mitochondrial dysfunction and involves oxidative stress through enhanced mitoROS generation. **(A)** Left: longitudinal quantitation of DUX4 and ZSCAN4-tdTomato containing nuclei in iDUX4^ZSCAN4-tdT^ reporter myoblasts at 2h intervals after DOX (62.5 ng/mL). Right: immunofluorescence micrographs of iDUX4^ZSCAN4-tdT^ reporter myoblasts before (0h), and 4h and 16h after exposure to DOX (62.5 ng/mL; green=DUX4, red=ZSCAN4-tdTomato, blue=nuclei; scale bar=100μm). **(B)** Normalised oxygen consumption rate (OCR) curves of iDUX4 control (CTRL) and DOX-induced (62.5 ng/mL DOX for 4h and 16h) myoblasts. **(C)** Respirometric assessment of mitochondrial respiration reveals reduction of basal, maximal and ATP-linked respiration in iDUX4 myoblasts 4h after DOX (62.5 ng/mL). **(D)** Normalised extracellular acidification rate (ECAR) curves of iDUX4 control (CTRL) and DOX-induced (62.5 ng/mL DOX for 4h and 16h) myoblasts. **(E)** ATP production rates demonstrating reduction of ATP in iDUX4 myoblasts 4h after DOX (62.5 ng/mL), mainly arising from defective oxidative phosphorylation (OXPHOS; mitoATP), while glycoATP is unaffected before 16h. **(F)** Assaying of Casp9 (red; onset 4h after DOX) and Casp3/7 activation (grey; onset 8h after DOX), measured at 2h intervals over 16h. **(G)** Assaying Annexin V as a marker of apoptosis in iDUX4 myoblasts after DOX (62.5 ng/mL), measured at 2h intervals over 16h. **(H)** Casp3/7 activation in iDUX4 myoblasts 8h after DOX (62.5 ng/mL) is prevented by a Casp9 inhibitor (Z-LEHD-FMK TFA; 10 μM). **(I)** Casp3/7 and Casp9 activation in iDUX4 myoblasts 8h after DOX (62.5 ng/mL) with non-targeted (Q10; 20 μM) or mitochondria-targeted (mitoQ10; 20μM) Coenzyme Q10. [n=3-6, data is mean ± s.d., where \**p*<0.05, \*\**p*<0.01, \*\*\**p*<0.001, \*\*\*\**p*<0.0001].

Crucially, DUX4-induced apoptosis was dependent on the intrinsic pathway, since inhibition of Casp9 with Z-LEHD-FMK TFA prevented DUX4-induced Casp3/7 activation (Fig. 3H). Furthermore, since oxidative stress is a well-established inducer of the mitochondrial apoptotic cascade [39], we next explored whether enhanced mitoROS generation from dysfunctional mitochondria contributes to Casp9 activation. We compared the ability of conventional (CoQ10) and mitochondria-targeted (mitoQ10) antioxidants to suppress Casp3/7 activation. Although Q10 supplementation of DUX4-expressing myoblasts slightly reduced Casp3/7 activation, the effect of mitoQ10 was much more pronounced as mitoQ10 also reduced Casp9 activation (Fig. 3I). Thus, mitoROS are involved in activation of pro-apoptotic mitochondrial signalling through Casp9.

### DUX4 drives mitochondrial oxidative stress and apoptosis through reverse electron transfer in the mitochondrial respiratory chain

Mechanisms of mitoROS generation in oxidative stress in FSHD and DUX4-expressing muscle cells remain unknown [32, 36, 40]. Since we identified redox regulation of Casp9 activation through mitoROS, we examined whether mitochondria, and the respiratory chain in particular, are the major ROS producing system. The mitochondrial respiratory chain can produce large amounts of ROS in skeletal muscles [41], but other mitochondrial and cytoplasmic mechanisms can contribute. Among these, NADPH oxidases (NOX; particularly NOX2 and 4 in skeletal muscle [42]), xanthine oxidase (XO) and uncoupled nitric oxide synthases (NOS) can produce significant amounts of ROS under pathological conditions [43]. Thus, we tested whether inhibition of NOX (Apocynin), XO (Allopurinol) or NOS (L-NAME) would reduce DUX4-induced ROS and Casp3/7 activation in iDUX4 myotubes (125 ng/mL DOX with inhibitor for 24h). No inhibitor reduced ROS levels or Casp3/7 activation, while NOS inhibition via L-NAME instead increased ROS and Casp3/7 activation (Fig. 4A), likely through removal of the natural ROS scavenger nitric oxide (NO·) [44]. Likewise, no inhibitor influenced ROS levels and Casp3/7 activation when DUX4 expression was induced in iDUX4 myoblasts (62.50 ng/mL DOX with inhibitor for 16h; Fig. S3A)

**Fig. 4:**
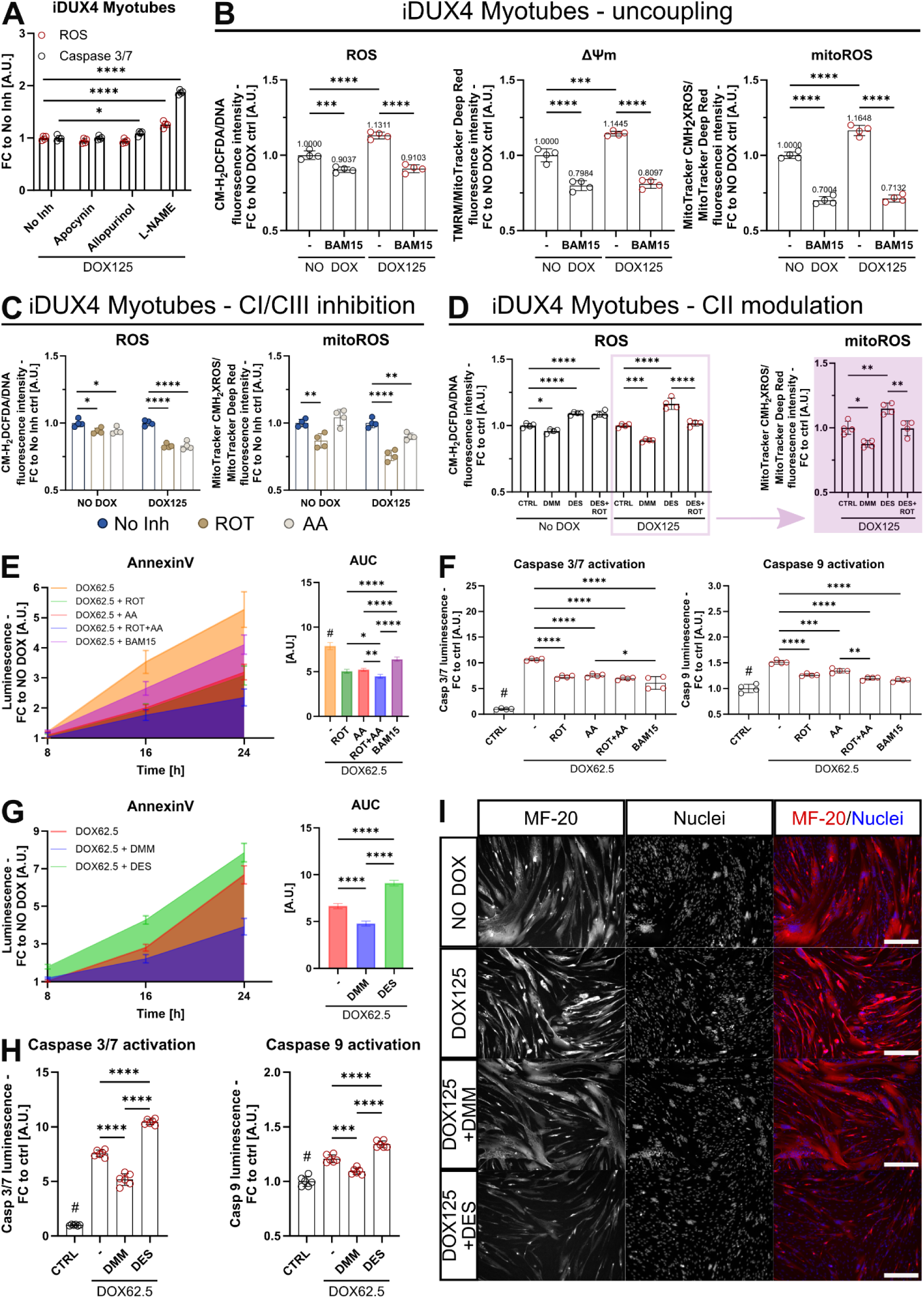
DUX4 causes oxidative stress-induced apoptosis via Caspase 9 through reverse electron transfer (RET)-ROS release from the respiratory chain in dysfunctional mitochondria. **(A)** ROS levels and Casp3/7 activation in iDUX4 myotubes after DUX4 induction with DOX (125 ng/mL) for 24h are not reduced upon inhibition of NOX (Apocynin 10 μM), XO (Allopurinol 10 μM) or NOS (L-NAME 5 mM). **(B)** ROS, ΔΨm and mitoROS are higher in iDUX4 myotubes after DUX4 induction with DOX (125 ng/mL) for 24h, with a greater reduction in DUX4-expressing myotubes compared to controls (NO DOX) after uncoupling of the mitochondrial respiratory chain with BAM15 (1 μM for 25 min). **(C)** ROS and mitoROS in iDUX4 myotubes after DOX (125 ng/mL) for 24h are disproportionally reduced in DUX4-expressing myotubes after Complex I (CI) inhibition with Rotenone (ROT, 0.5 μM/25 min) or Complex III (CIII) inhibition with Antimycin A (AA, 0.5 μM/25 min). **(D)** ROS levels in iDUX4 myotubes after DUX4 induction with DOX (125 ng/mL) for 24h with subsequent modulation of Complex II (CII)-linked respiration through the competitive CII inhibitor DMM (10 mM/25 min), the CII substrate DES (5 mM/25 min), or DES (5 mM for 25 min) and the CI inhibitor Rotenone (ROT, 0.5 μM for 25 min). Modulation of CII-linked respiration after DUX4 expression affects ROS by gauging mitoROS levels from the respiratory chain (purple insert). **(E)** Assaying Annexin V and AUC quantitation in iDUX4 myoblasts after DOX (62.5 ng/mL) over 24h with ROT (0.5 μM), AA (0.5 μM), ROT+AA (0.5 μM each) or BAM15 (1 μM) reveals reduced DUX4-induced toxicity when mitoROS release from the respiratory chain is suppressed. **(F)** Reduced Casp3/7 and Casp9 activation in iDUX4 myoblasts after DOX (62.5 ng/mL) over 8h (Casp9) or 24h (Casp3/7) in with ROT (0.5 μM), AA (0.5 μM), ROT+AA (0.5 μM each) or BAM15 (1 μM). **(G)** Assaying Annexin V and AUC quantitation in iDUX4 myoblasts after DOX (62.5 ng/mL) over 24h with competitive CII inhibitor DMM (10mM) or CII substrate DES (5mM). Reduction of CII-linked respiration (DMM) reduces, while induction of CII-linked respiration (DES) increases, DUX4-induced apoptosis. **(H)** Modulation of Casp3/7 and Casp9 activation in iDUX4 myoblasts after DOX (62.5 ng/mL) over 8h (Casp9) or 24h (Casp3/7) with DMM or DES. **(I)** Immunofluorescence of iDUX4 myotubes 36h after exposure to DOX (125 ng/mL) with concomitant administration of DMM (10 mM) or DES (5 mM), showing rescue (DMM) or aggravation (DES) of the hypotrophic myotube phenotype triggered by DUX4 [red=myosin heavy chain (MF-20), blue=nuclei; scale bar=200 μm]. [n=4, data is mean ± s.d., where \**p*<0.05, \*\**p*<0.01, \*\*\**p*<0.001, \*\*\*\**p*<0.0001; # denotes statistical significance from all other samples and untreated DOX62.5 control (-) **(E)** or non-DOX induced control (CTRL) **(F and H)**].

We therefore investigated mitochondrial respiratory chain involvement in ROS generation. DUX4 increases (mito)ROS levels in myotubes, driven by elevated ΔΨm, which could effectively be reduced through dissipation of ΔΨm by the mitochondria-specific uncoupler BAM15 (Fig. 4B). Of note, uncoupling of the respiratory chain reduced (mito)ROS levels in both controls and DOX-treated iDUX4 myotubes, but to a greater extent when DUX4 was expressed, up to the level of uncoupled controls. Since complex I and III in the respiratory chain are known ROS producers [45], we examined effects of short-term inhibition of complex I with Rotenone (ROT), or complex III with Antimycin A (AA). Both ROT and AA moderately reduced ROS levels in non-induced iDUX4 control myotubes, but this was more significant when DUX4 was induced (Fig. 4C). Interestingly, complex III inhibition reduced mitoROS levels in DUX4-expressing myotubes only, while complex I inhibition reduced mitoROS levels in both non-induced and, to a greater extent, DUX4-induced myotubes.

Under forward electron transfer (FET) through the mitochondrial ETC, inhibition of complex I and/or III tends to increase mitoROS generation [46]. Thus, we hypothesised that mitoROS production in response to DUX4 induction could be driven by RET, where electrons flow ‘backwards’ through the ETC, leading to production of large amounts of ROS, especially by complex I. Since ΔΨm increases after DUX4 induction in iDUX4 myotubes prior to cell death (Fig. 4B) and high ΔΨm favours RET, we measured ROS under modulation of complex II activity. The competitive complex II inhibitor dimethyl malonate (DMM) reduced ROS more efficiently in iDUX4 myotubes, while stimulation of complex II-linked respiration with diethyl succinate (DES) increased ROS disproportionally after DUX4 induction compared to non-induced controls (Fig. 4D). Of note, complex I inhibition with ROT diminished DES-induced ROS increase in DUX4-expressing myotubes, but not in controls, and changes in ROS levels were driven by concomitant mitoROS modulation (Fig. 4D purple insert). Observations in iDUX4 myotubes were consistent with DUX4-induced changes in iDUX4 myoblasts (Fig. S3B, C, E) and show that mitoROS generation through RET is an oxidative stress mechanism peculiar to muscle cells exposed to DUX4.

Since DUX4-induced apoptosis is, at least in part, regulated by mitoROS (Fig. 3I), we investigated effects of uncoupling (BAM15) or complex I/III inhibition (ROT, AA). All compounds significantly reduced apoptosis in iDUX4 myoblasts (Fig. 4E) through reduction of Casp3/7 and 9 activation (Fig. 4F). Dose-response testing with simultaneous inhibition of complex I (ROT) and III (AA) revealed a synergistic effect of dual inhibition (Fig. S3D), as assessed with Highest Single Agent (HAS) synergy scoring. Importantly, neither uncoupling nor complex I/III inhibition affected DUX4 itself but complex I inhibition showed a moderate reduction of DUX4-mediated activation of ZSCAN4 in iDUX4^ZSCAN4-tdT^ myoblasts (Fig. S3F).

We also measured apoptosis in iDUX4 myoblasts under modulation of complex II with DMM or DES to gauge RET-ROS formation. Complex II inhibition with DMM reduced DUX4-induced apoptosis by suppressing Casp3/7 and 9 activation, while stimulation with DES had the opposite effect and aggravated DUX4-induced cell death (Fig. 4G, H). A similar response to RET-ROS inhibition/stimulation was observed in iDUX4 myotubes, where DMM inhibition rescued DUX4-induced myotube hypotrophy, while DES stimulation aggravated this phenotype, as assessed by immunolabeling for myosin heavy chain (Fig. 4I).

### RET-ROS release from the respiratory chain is an oxidative stress mechanism in FSHD mitochondria and induces mitochondrial apoptosis through Caspase 9

We also investigated mitochondrial respiratory chain involvement in our three independent FSHD patient-derived muscle models. Both FSHD myoblasts and myotubes (54-6, K4 and 16U) had higher steady-state activation of Casp3/7 and 9 compared to matched controls (54-12, K8, 16A; Fig. S4A), showing higher baseline activity of the intrinsic apoptotic pathway in FSHD. Consistent with the iDUX4 model, NOX, XO and NOS did not contribute to ROS generation and Casp3/7 activation in FSHD myotubes, while NOS inhibition using L-NAME increased ROS levels and Casp3/7 activation (Fig. 5A). Dissipation of ΔΨm by the uncoupler BAM15 diminished elevated ROS observed in FSHD myotubes by reducing mitoROS generation (Fig. 5B). Notably, inhibition of complex I with ROT and complex III with AA reduced (mito)ROS in FSHD myotubes but not controls (Fig. 5C). These effects were consistent between our three FSHD cellular models, and only observed in FSHD myotubes (Fig. S4B, C).

**Fig. 5:**
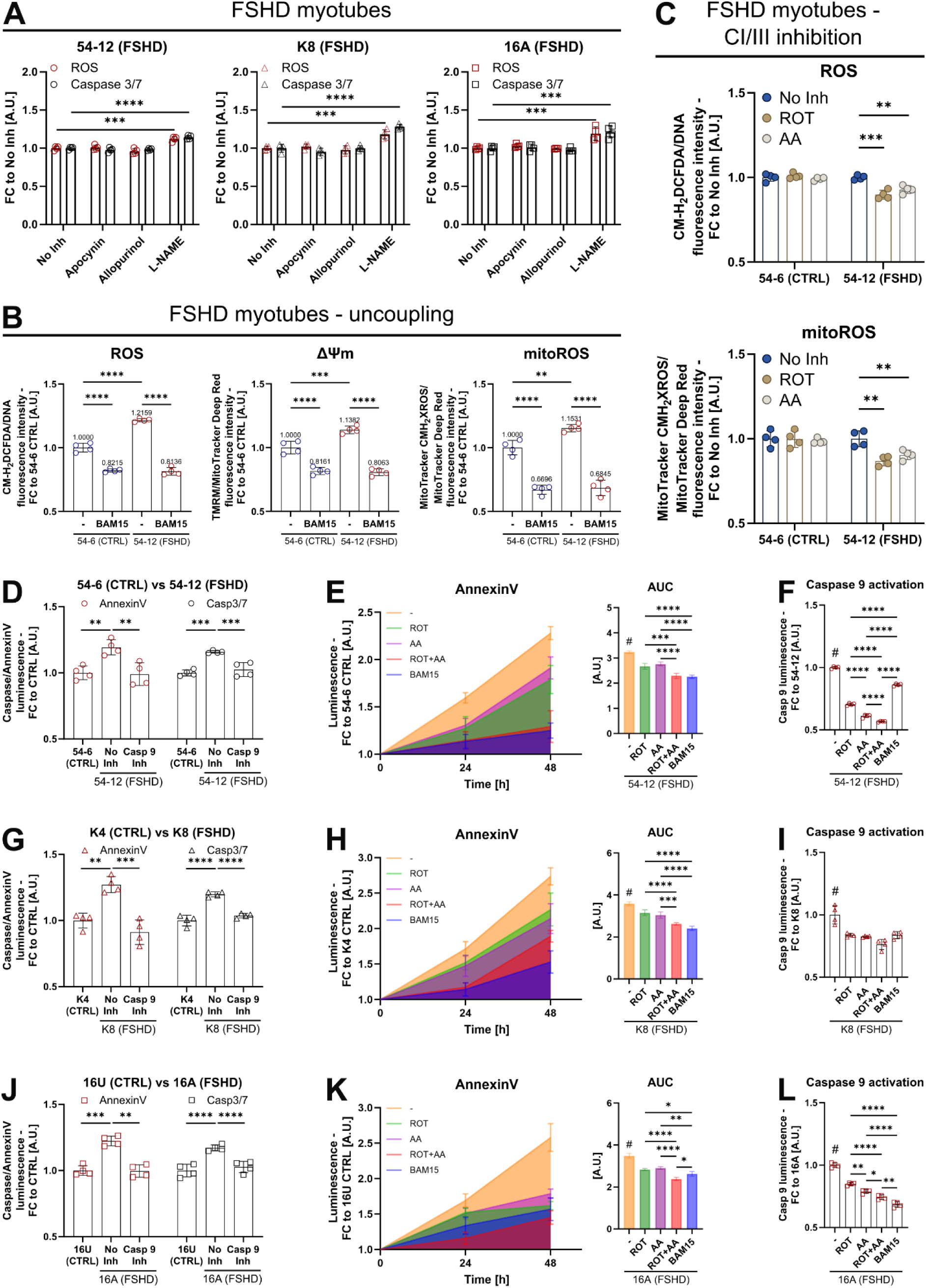
Dysfunctional mitochondria are the main ROS producing system in FSHD muscle cells and trigger apoptosis in a Caspase 9-dependent manner. **(A)** ROS and Casp3/7 activation in FSHD myotubes (54-12, K8, 16A) are not reduced when NOX (Apocynin 10 μM), XO (Allopurinol 10 μM) or NOS (L-NAME 5 mM) are inhibited for 24h. **(B)** ROS, ΔΨm and mitoROS are higher in FSHD myotubes (54-12), with a greater reduction in FSHD compared to control myotubes (54-6) after uncoupling the mitochondrial respiratory chain with BAM15 (1 μM/25 min). **(C)** ROS and mitoROS are reduced in FSHD myotubes (54-12) after CI inhibition with Rotenone (ROT, 0.5 μM/25 min) or CIII inhibition with Antimycin A (AA, 0.5 μM/25 min), but not in controls (54-6). **(D, G and J)** Elevated baseline apoptosis and Casp3/7 activation in FSHD myoblasts (54-12, K8 and 16A) is reduced to control (54-6, K4 and 16U) levels through inhibition of Casp9 (Z-LEHD-FMK TFA; 10 μM/24h). **(E, H and K)** Assaying of Annexin V over 48h in FSHD myoblasts (54-12, K8 and 16A) with ROT (0.5 μM), AA (0.5 μM), ROT+AA (0.5 μM each) or BAM15 (1 μM) and AUC quantitation demonstrates reduced apoptosis when mitoROS release from the respiratory chain is suppressed in FSHD myoblasts. **(F, I and L)** Reduced Casp9 activation in FSHD myoblasts (54-12, K8 and 16A) after administration of ROT (0.5 μM), AA (0.5 μM), ROT+AA (0.5 μM each) or BAM15 (1 μM) for 48h. [n=4, data is mean ± s.d., where \**p*<0.05, \*\**p*<0.01, \*\*\**p*<0.001, \*\*\*\**p*<0.0001; # denotes significant difference of untreated control (-) compared to all other samples].

Higher apoptosis (Annexin V) and Casp3/7 activation in FSHD myoblasts (Fig 5 D-L) was rescued by Casp9 inhibition with Z-LEHD-FMK TFA (Fig. 5D, G, J), and uncoupling or complex I/III inhibition reduced apoptosis by suppressing Casp9 activation, with an additive effect of combined complex I and III inhibition (Fig. 5.E, F, H, I, K, L).

RET-ROS generation was evident in FSHD myotubes, in line with the iDUX4 model. Complex II inhibition with DMM did not affect ROS levels in controls (54-6, K4, 16U), but significantly and consistently reduced ROS levels in FSHD myotubes (54-12, K8, 16A). Stimulation of complex II through DES increased ROS levels in control and FSHD myotubes alike, but this was uniquely diminished by simultaneous complex I inhibition with ROT in FSHD myotubes (Fig. 6A, C, E). Changes in ROS levels in response to complex II modulation and complex I inhibition under DES-stimulated RET-ROS formation were driven by mitoROS production in FSHD mitochondria (Fig. 6A, C, E purple inserts). RET-ROS were involved in FSHD apoptosis by modulating mitochondrial apoptosis, with apoptosis and Casp9 activation reduced after DMM treatment, while DES increased FSHD apoptosis (Fig. 6B, D, F). Likewise, myogenesis was uniquely affected by RET-ROS in FSHD cells: while DMM and DES did not affect myotube formation in controls, DMM inhibition rescued the hypotrophic FSHD myotube phenotype, with DES further aggravating this phenotype, as assessed by immunolabeling for myosin heavy chain (Fig. 6G).

**Fig. 6:**
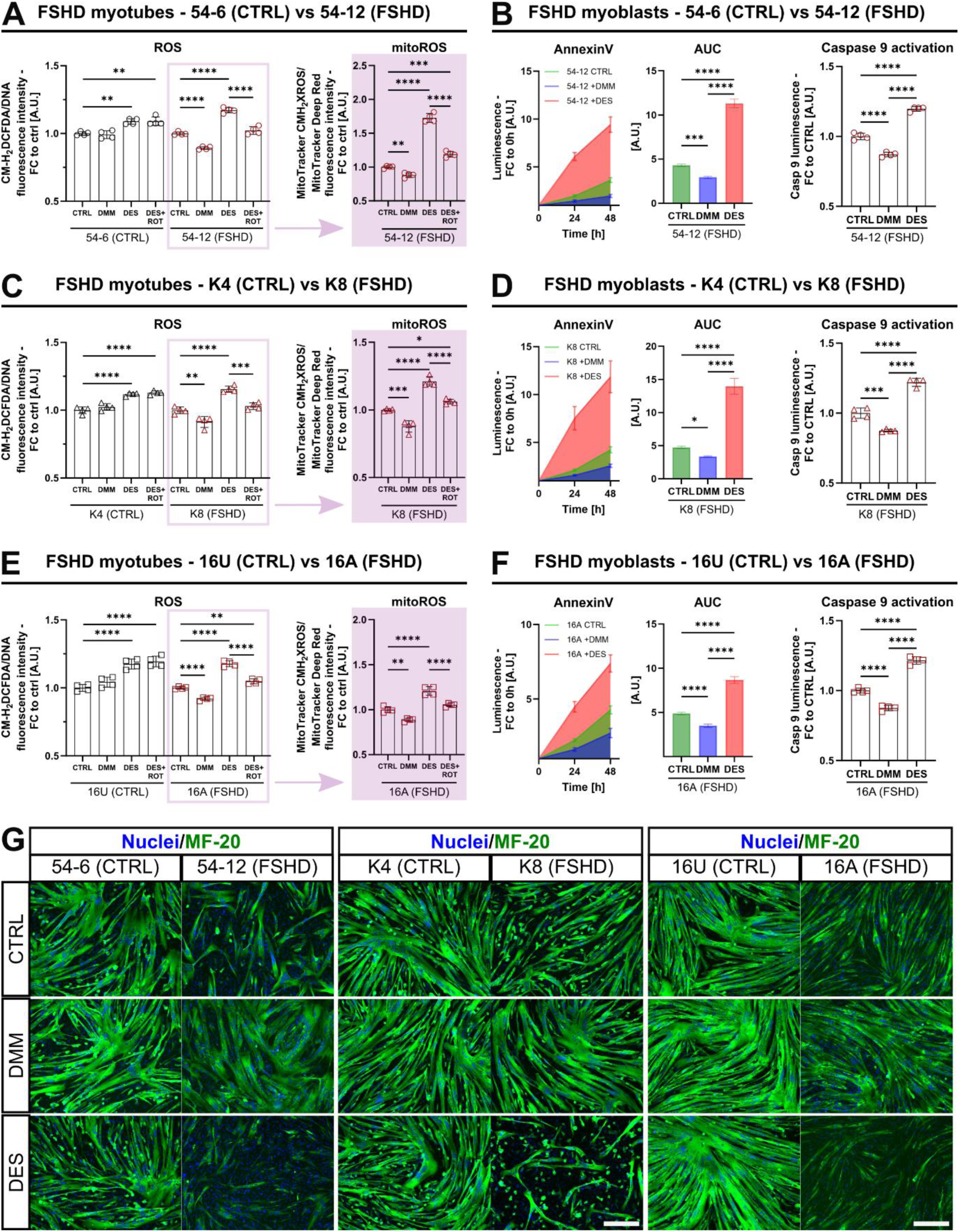
RET-ROS release from the respiratory chain is a unique feature of FSHD mitochondria and induces mitochondrial apoptosis and myotube hypotrophy in FSHD. **(A, C and E)** ROS levels in control 54-6, K4 and 16U and FSHD 54-12, K8 and 16A myotubes after modulation of complex II (CII)-linked respiration through administration of DMM (10 mM/25 min), DES (5 mM/25 min), or both DES (5 mM/25 min) and Rotenone (ROT, 0.5 μM/25 min). Modulation of CII-linked respiration affects ROS uniquely in FSHD myotubes by gauging mitoROS from the respiratory chain (purple insert). **(B, D and F)** Left: Assaying of Annexin V and AUC quantitation in FSHD 54-12, K8 and 16A myoblasts over 48h with DMM (10 mM) or DES (5 mM). Reduction of CII-linked respiration (DMM) reduced, while induction of CII-linked respiration (DES) increased, FSHD myoblast apoptosis. Right: Casp9 activation in FSHD 54-12, K8 and 16A myoblasts is reduced by DMM (10 mM/8h) and enhanced by DES (5 mM/8h). **(G)** Immunofluorescence of FSHD myotubes (54-12, K8, 16A) differentiated with DMM (10 mM) or DES (5 mM), alongside controls (54-6, K4, 16U). DMM rescues, while DES aggravates, the hypotrophic myotube phenotype in FSHD myotubes, but does not affect controls [green=myosin heavy chain (MF-20), blue=nuclei; scale bar=200μm]. [n=4, data is mean ± s.d., where \**p*<0.05, \*\**p*<0.01, \*\*\**p*<0.001, \*\*\*\**p*<0.0001].

### Transcriptomic analysis indicates perturbed mitochondrial respiratory chain function and apoptosis as features of FSHD patient muscle

Having identified RET as a mitochondrial oxidative stress mechanism crucial for DUX4 toxicity in muscle cells, we next assessed whether FSHD patient muscles show evidence of this mechanism. We performed enrichment Over-Representation Analysis (ORA) of HALLMARK gene sets on previously reported RNA-seq data from magnetic resonance imaging (MRI)-informed FSHD muscle biopsies for transcriptomic changes specific to metabolic pathways, OXPHOS, respiratory chain function and apoptosis. Wong et al. [47] stratified 36 FSHD patients into four groups based on clinical/histopathological criteria and DUX4 target gene expression (G1: lowest to G4: highest disease severity), alongside 9 unaffected controls (Fig. 7A and Supplementary File 1). Given that biopsies were taken under MRI-guidance to visualise oedema as footprints of active muscle pathology (T2-weighted Short Tau Inversion Recovery sequences), genes involved in ‘Inflammatory Response’ were upregulated irrespective of FSHD disease severity (Fig. 7B). Genes involved in apoptosis, p53 pathway and hypoxia were increased in G1-4, as was Interleukin 6-Janus kinase-Signal Transducer and Activator of Transcription (IL6-JAK-STAT) signalling - a convergent pathway on the interface of inflammation and hypoxic stress. Metabolic pathways with mitochondrial involvement (OXPHOS, fatty acid metabolism, heme metabolism, bile acid metabolism) were downregulated in FSHD patients with high disease severity (G4), with OXPHOS the biggest and most significantly enriched downregulated HALLMARK gene set (Fig. 7B). The latter is consistent with a recent transcriptomic meta-analysis of multiple FSHD cell- or patient-derived RNA-seq and microarray datasets identifying many OXPHOS/ETC genes among the most repressed genes in FSHD [28]. Notably, genes involved in providing oxidation substrates for mitochondrial respiratory chain entry sites at complex I and II (Fig. 7C, red insert) and in complex I-linked mitochondrial electron transfer showed downregulation at the highest disease severity G4 (Fig. 7C, black insert). This is in line with our data demonstrating that DUX4 affects predominantly complex 1-linked respiration in muscle cells [32], and supports our observation that complex I drives oxidative stress through RET.

**Fig. 7:**
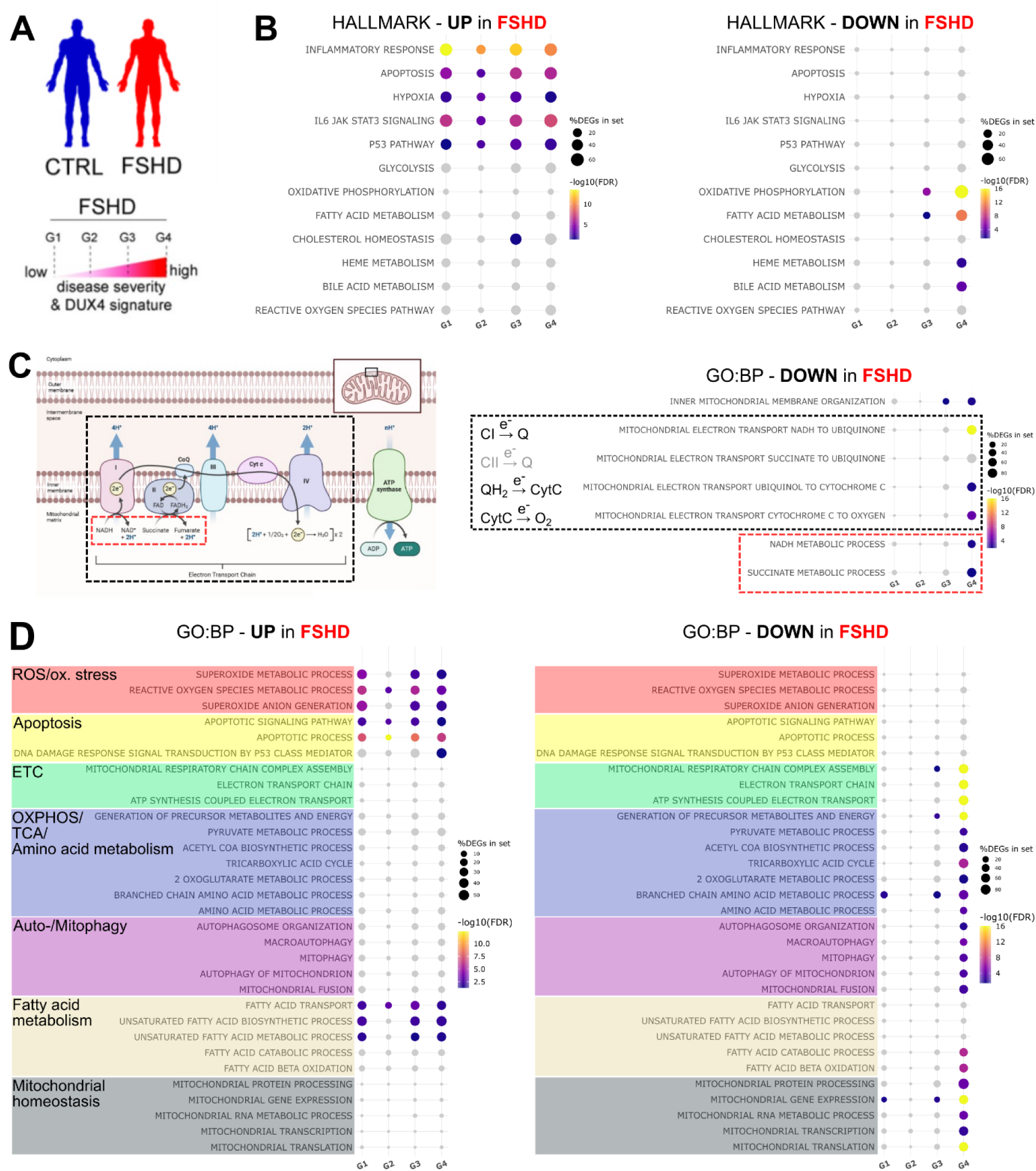
Transcriptional repression of mitochondrial redox-linked metabolic pathways is a feature of FSHD muscle. **(A)** HALLMARK enrichment over-representation analysis (ORA) in muscle biopsies from 36 FSHD patients with varying degrees of disease severity (Ctrl: healthy control individuals; G1: lowest disease severity score; G4: highest disease severity score based on clinical assessment, histopathology scoring and DUX4 target gene expression) [47]. **(B)** Pathways involved in mitochondrial metabolism (OXPHOS and fatty acid metabolism) are downregulated in FSHD patients with high severity (G3 and G4), whereas pathways involved in inflammation, apoptosis and molecular response to hypoxia are upregulated across all 4 disease severity groups (grey circles indicate non-significant enrichment of respective gene sets based on −log10(FDR). **(C)** Downregulation of genes specifically involved in mitochondrial electron transport through the respiratory chain (red box: complex I and II oxidation substrates NADH and succinate, respectively; black box: stages of electron transport through respiratory chain complexes I, II, III and IV) in muscle biopsies from FSHD patients with high disease severity (G4). Complex I-linked respiration is perturbed at all stages through the ETC in G4. Scheme of the mitochondrial respiratory chain was created with BioRender.com. **(D)** Enrichment ORA of Gene Ontology:Biological Processes (GO:BP) gene sets relating to oxidative stress (orange), apoptosis (yellow), ETC (green), OXPHOS/TCA/amino acid metabolism (blue), auto- and mitophagy (purple), fatty acid metabolism (beige) and mitochondrial homeostasis (grey). ROS metabolism, apoptosis and anabolic fatty acid metabolism are the major upregulated metabolic clusters in FSHD patient muscle across disease severity groups. Repression of pathways related to (oxidative) mitochondrial metabolism (ETC, OXPHOS, TCA, amino acid metabolism, fatty acid breakdown and mitochondrial transcription/translation) marks FSHD patient muscle with high severity scores (G4). Data shown as pairwise contrasts of Ctrl versus G1-G4 disease severity groups.

To further investigate transcriptional changes pertaining to perturbation of oxidative (mitochondrial) metabolism, we performed enrichment ORA of Gene Ontology:Biological Process (GO:BP) gene sets relating to mitochondrial function, turnover and homeostasis (Fig. 7D and Supplementary File 2). In general, mitochondrial metabolic GO:BP gene sets dominated the most significantly repressed pathways, with consistent upregulation of genes involved in ROS metabolism (orange box) and apoptosis (yellow box) across many disease severity groups. Conversely, genes involved in the ETC (green box), OXPHOS/TCA/amino acid metabolism (blue box), auto- and mitophagy (purple box) and mitochondrial homeostasis (grey box) were repressed in FSHD patients with high disease severity (G4), indicating widespread metabolic/mitochondrial stress in FSHD muscles. Interestingly, cytoplasmic-based anabolic fatty acid metabolism was upregulated across many disease severity groups, while mitochondrial-based fatty acid catabolism through β-oxidation was repressed in G4, supporting mitochondrial dysfunction as a driver of perturbed fatty acid breakdown which may contribute to the energy deficit observed in FSHD muscle fibres [27].

### The mitochondrial respiratory chain is required for DUX4-induced apoptosis in human muscle cells

Our findings indicate a central involvement of the mitochondrial respiratory chain and RET as a ROS producer and pro-apoptotic signalling entity in response to DUX4 activation. Since RET-ROS originate from the ETC, we generated ‘ρ^0^-like’ iDUX4 myoblast lines with varying degrees of OXPHOS impairment (Fig. 8A) to further assess interaction between DUX4-induced downstream events and the mitochondrial respiratory chain. Through titration of ethidium bromide (EtBr) for two-weeks we obtained iDUX4-ρ^0^ lines with moderate (iDUX4-ρ^0+^), strong (iDUX4-ρ^0++^) or full (iDUX4-ρ^0+++^) OXPHOS impairment, as determined by respirometry (Fig. 8B). Fully OXPHOS deficient iDUX4-ρ^0+++^ myoblasts were unable to produce ATP through OXPHOS (Fig. S5A, B) but compensated by enhancing glycolytic ATP production (Fig. 8B). Further, iDUX4-ρ^0+++^ did not show overt changes in mitochondrial morphology but exhibited ∼50% mtDNA depletion, drastically reduced ΔΨm and mitoROS, and uridine auxotrophy (Fig. S5C-E). Lack of mitochondrial respiratory chain function in iDUX4-ρ^0+++^ myoblasts did not affect expression of the *DUX4* transgene after DOX administration (Fig. 8C), although post-induction levels of DUX4 target genes *ZSCAN4*, *PRAMEF1* and *TRIM43* were lower (Fig. 8D). In iDUX4-ρ^0+++^ myoblasts devoid of functional OXPHOS, DUX4 induction was unable to trigger increased ΔΨm, mitoROS and Casp9 activation (Fig. 8E, F), unlike the situation in unmodified iDUX4 controls (Fig. 4B, F and Fig. S3B). DUX4-induced apoptosis and Casp3/7 activation after long term (36h) induction with DOX correlated inversely with the level of OXPHOS impairment, and, crucially, cell death was not observed in iDUX4-ρ^0+++^ myoblasts with full OXPHOS deficiency (Fig. 8G). However, iDUX4-ρ^0+++^ myoblasts readily underwent apoptosis in response to staurosporine (STSP) at levels comparable to unmodified iDUX4 controls (Fig. 8H). Since iDUX4-ρ^0+++^ cannot produce mitoROS through RET-ROS to activate Casp9, we attribute this protection against DUX4-mediated cell death to the lack of a functioning mitochondrial respiratory chain.

**Fig. 8:**
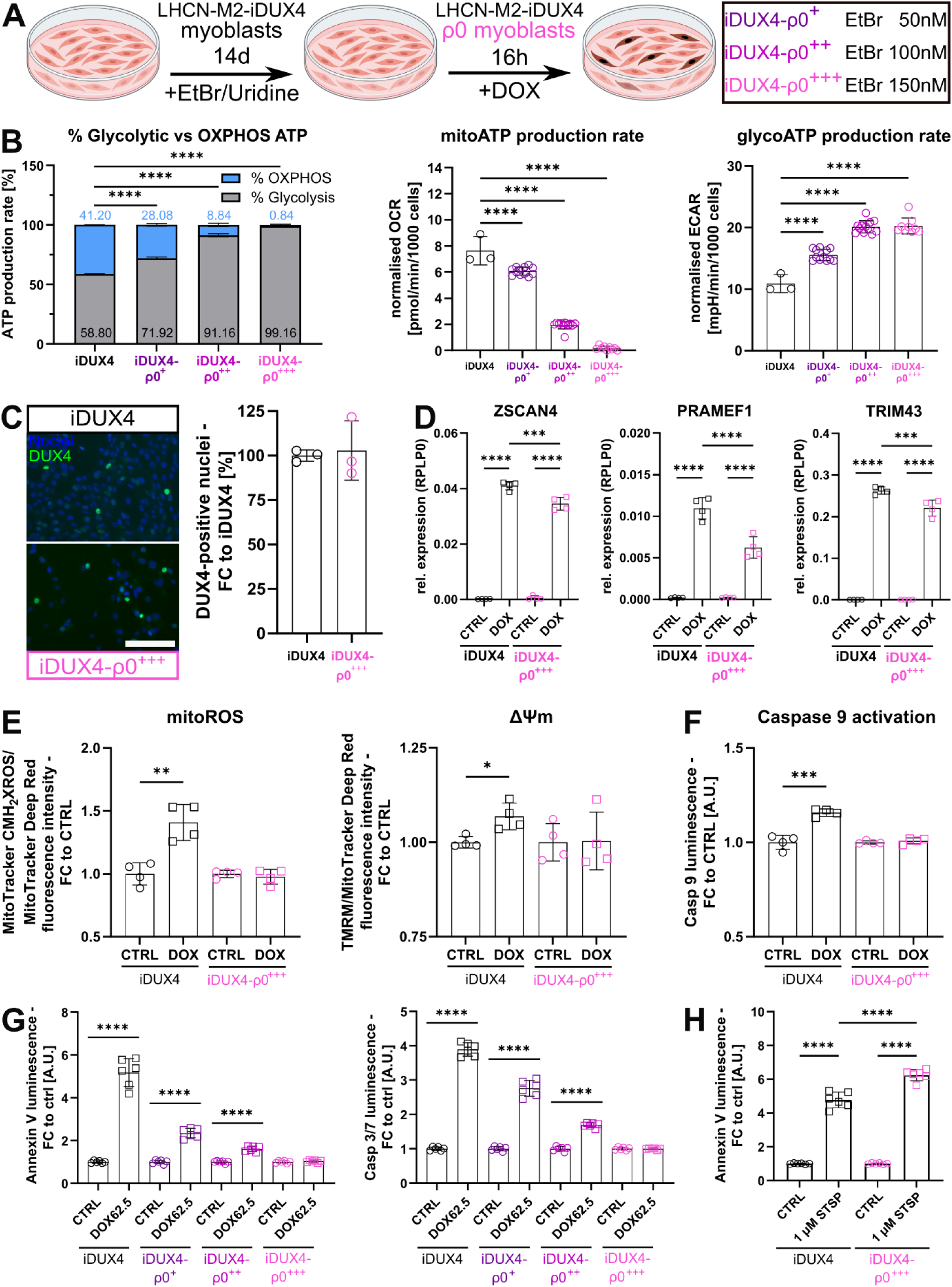
Mitochondrial respiratory chain function is required for DUX4 toxicity in human muscle cells. **(A)** Workflow for generation of ρ^0^-like iDUX4 myoblasts: cells were treated with ethidium bromide (50, 100 or 150 nM EtBr) for two weeks, yielding ρ^0^-like iDUX4 cells with varying degrees of OXPHOS impairment (iDUX4-ρ^0+^: mild OXPHOS reduction, iDUX4-ρ^0++^: severe OXPHOS reduction and iDUX4-ρ^0+++^: full OXPHOS inhibition). Scheme was created with BioRender.com. **(B)** Percentage of OXPHOS versus glycolytic ATP production rate in iDUX4-ρ^0^ myoblasts, with mitoATP and glycoATP production rates indicated, as determined by Seahorse respirometry. **(C)** Immunofluorescence (green=DUX4, blue=nuclei; scale bar=100μm) with percentage of DUX4-positive nuclei of unmodified iDUX4 and iDUX4- ρ^0+++^ myoblasts after DOX (62.5 ng/mL) for 16h. **(D)** Expression levels of DUX4 target genes *ZSCAN4*, *PRAMEF1* and *TRIM43*, as assessed by RT-qPCR (relative to housekeeper *RPLP0*). **(E)** DUX4-induced increases in mitoROS and ΔΨm in unmodified iDUX4 cells are not observed in iDUX4- ρ^0+++^ myoblasts after DOX (62.5 ng/mL) for 16h. **(F)** DUX4-induced Casp9 activation in unmodified iDUX4 myoblasts does not occur in iDUX4- ρ^0+++^ cells after DOX (62.5 ng/mL) for 8h. **(G)** Annexin V and Casp3/7 activation in iDUX4, iDUX4-ρ^0+^, iDUX4-ρ^0++^ and iDUX4-ρ^0+++^ myoblasts after DOX (62.5 ng/mL) for 36h shows inverse correlation between DUX4-induced apoptosis and mitochondrial respiratory chain impairment. DUX4 is unable to induce apoptosis in fully OXPHOS-deficient iDUX4-ρ^0+++^ myoblasts. **(H)** Both non-DUX4-induced iDUX4 and iDUX4-ρ^0+++^ myoblasts undergo apoptosis (Annexin V) after exposure to the apoptosis inducer staurosporine (STSP; 1 μM) for 48h. [n=3-12, data is mean ± s.d., where \**p*<0.05, \*\**p*<0.01, \*\*\**p*<0.001, \*\*\*\**p*<0.0001].

## DISCUSSION

We show that mitochondria, particularly through their respiratory chain, are directly involved in DUX4-mediated damage in FSHD. RET-driven mitoROS generation from complex I is a major mechanism of DUX4-induced oxidative stress generation. These RET-ROS are involved in activation of the intrinsic (mitochondrial) apoptotic pathway through Casp9. Importantly, DUX4 requires mitochondrial respiratory chain function to exert toxicity. We also demonstrate that RET is a mitochondrial oxidative stress mechanism specifically present in FSHD cells compared to controls, as mitoROS in control cells were not reduced by complex II inhibitor DMM or complex I inhibitor ROT. RET-ROS release from complex I as the main driver of mitoROS formation is triggered by DUX4 induction independent of muscle differentiation. Crucially, inhibition of complex II-linked respiration with DMM rescues abnormal FSHD phenotypes, reducing or normalising mitoROS generation, apoptosis and myotube hypotrophy. Integrated use of DUX4-inducible and FSHD cell lines indicates that RET is a direct consequence of *DUX4* misexpression in FSHD models, linking this mitochondrial pathomechanism directly to epigenetic de-repression of *DUX4*.

RET is a potent mitoROS production mechanisms and became widely known as the driver of oxidative damage in cardiac ischemia-reperfusion injury [48, 49], where RET-ROS production is caused by a highly reduced CoQ pool and a large proton motive force (Δp and ΔΨm) across the inner mitochondrial membrane. Together, these force electrons back through the ETC to complex I, resulting in reduction of NAD^+^ to NADH and a dramatic increase in complex I-linked superoxide (O^2·-^) production [50]. RET at complex I is now an established (patho)physiological redox mechanism [51–53] and strategies to prevent oxidative damage from excessive RET-ROS generation in ischemia-reperfusion include manipulating ΔΨm, complex I- and/or complex II-linked respiration [54].

Here, we also employed uncoupling (BAM15), complex I inhibition (ROT) and complex II inhibition (DMM) strategies, and showed that DMM is efficient in reducing RET-ROS and mitochondrial apoptosis in iDUX4 (Fig. 4) and FSHD cell lines (Fig. 5, 6). RET is a major ROS generating mechanism in FSHD as inhibition of other ROS producers had no protective effects against DUX4-induced cell death. While 70% of total ROS is attributed to complex I and III, ∼30% comes from NOX inhibition in skeletal muscle [45]. However, *DUX4* expression in iDUX4-ρ^0^ cells reveals that DUX4-induced apoptosis requires respiratory chain function (Fig. 7), making our finding that complex III inhibition reduces mitoROS and apoptosis surprising. AA inhibits electron flow through the Q_i_ site in complex III, thus inhibiting the Q cycle and stimulating ROS generation from the Q_o_ site [55, 56]. Under conditions favouring RET, where the QH_2_/Q ratio is high, Q_i_ site inhibition would feed into overreduction of the Q pool and enhance RET-ROS release. Cells were with respiratory chain inhibitors for only 25 min. Since RET is driven by high ΔΨm, reduction in ROS levels by AA must be driven by a collapse of ΔΨm, an immediate consequence of reduced electron flow from the Q pool via complex III to cytochrome C (before AA stimulates ROS production from the Q_o_ site in complex III upon longer exposure) [57].

Transcriptomic changes indicative of oxidative metabolic impairment (specifically OXPHOS and fatty acid metabolism) can be identified in FSHD muscle (Fig. 7). However, mitochondrial respiratory chain impairment in FSHD muscles suggests a central role for DUX4-driven transcriptional perturbation of the ETC in the aetiology of mitochondrial dysfunction and oxidative stress in FSHD. Respirometry in FSHD myotubes demonstrated functional OXPHOS deficits, evident as global reduction of ATP-linked respiration and altered balance between glycolytic and oxidative ATP generation (Fig. 1). This is consistent with the demonstration that FSHD myofibres *ex vivo* exhibit reduced ATP generation through OXPHOS [27]. We also identified a glycolytic ATP production deficit in FSHD cellular models. This aligns with a recent proteomic analysis in FSHD muscle biopsies that reveals glycolytic deficits, alongside perturbed OXPHOS, TCA cycle and fatty acid β-oxidation [29], in contrast to our transcriptomic analyses which suggested no glycolytic perturbation. Interestingly, the same study also showed that NAD(P)^+^/NAD(P)H ratios were drastically reduced with increasing disease severity in FSHD muscles, indicating RET and consequential NAD^+^ reduction to NADH through complex I. Such discrepancy between transcriptomic and proteomic data underscores the importance of investigating metabolic changes also at the effector level.

Little is known about metabolic coupling of glycolysis and OXPHOS in FSHD, particularly what causes glycolytic perturbations, and whether the primary driver of DUX4-induced metabolic stress is glycolysis or OXPHOS. These pathways are tightly interconnected since glycolysis influences OXPHOS through provision of pyruvate, which fuels the TCA cycle after its conversion to acetyl-CoA, and in turn, glycolysis requires NAD^+^, which is regenerated by NADH oxidation through OXPHOS [58]. DUX4 misexpression in myotubes primarily affects OXPHOS (Fig. 2), as the relative increases in glycolytic ATP rates where predominantly propelled by deficits in mitoATP production. In addition, kinetic profiling of ATP production demonstrated an immediate reduction of mitoATP upon DUX4 target gene activation, while glycoATP was affected later, and only moderately (Fig. 3). DUX4-induced OXPHOS inhibition as the main metabolic insult in FSHD is further supported by moderate glycolytic impairment at the proteomic level, but pronounced OXPHOS deficits in FSHD muscles with mild disease severity [29], and transcriptional repression of several respiratory chain complex subunits [28]. However, glucose metabolism and OXPHOS are both sensitive to regulation by ROS [59], and the possibility of DUX4 interfering with the response to hypoxia through perturbation of hypoxia-inducible factor 1α (HIF1α) signalling [32, 60–62] adds complexity. Since HIF1α stabilisation and subsequent metabolic adaptation can be caused by mitoROS [63] and/or succinate build-up [64], we speculate that DUX4-HIF1α crosstalk could be also mediated by RET, which in turn could trigger a persistent pro-inflammatory muscle state by chronic activation of the IL6-JAK-STAT3 axis.

Exact mechanisms by which DUX4 causes mitochondrial functional changes, especially given the notoriously low DUX4 levels in FSHD patients, still requires elucidation. Onset of DUX4-induced OXPHOS defects coincides with DUX4 target gene activation (Fig. 3). Very low DUX4 transcript and protein levels in FSHD cellular models suggest an epigenetic regulation of OXPHOS genes. In addition, DUX4 activates some transcriptional targets indirectly through ROS [36]. Our iDUX4-ρ^0^ cells show that the respiratory chain modulates activation of some DUX4 target genes through mitoROS, as ZSCAN4, PRAMEF1 and TRIM43 were expressed at lower levels upon *DUX4* expression in iDUX4-ρ^0^ cells than in unmodified iDUX4 (Fig. 8). RET-ROS contributed to this modulation, as ZSCAN4 activation upon DUX4 induction in iDUX4^ZSCAN4-tdT^ cells was reduced by complex I inhibition with ROT, but not DUX4 levels (Fig. S3).

In conclusion, we identified a major DUX4-induced oxidative stress mechanism, whereby dysfunctional mitochondria produce mitoROS through RET in the ETC, potentially playing a relevant role in FSHD pathophysiology. DUX4 myotoxicity significantly relies on this mechanism, as pharmacological inhibition of RET-ROS generation reduces DUX4-induced apoptosis, and DUX4 toxicity was not observed in OXPHOS deficient muscle cells. Importantly, RET-ROS formation can be pharmacologically targeted, for example through uncouplers or specific respiratory chain complex inhibitors, opening the way for the modulation of the mitochondrial respiratory chain as a potential treatment for FSHD.

## MATERIALS AND METHODS

### Reagents

Doxycycline (D5207), Blasticidine S (R21001), BAM15 (SML1760), Rotenone (ROT; R8875), Antimycin A (AA; A8674), CoenzymeQ10 (Q10; C95380, Apocynin (178385), Allopurinol (A8003), diethyl succinate (DES; 112402), dimethyl malonate (DMM; 136441), ethidium bromide (EtBr; E1510) and Uridine (U3750) were from Sigma-Aldrich (St. Louis, MO, USA). Oligomycin A (ab1434230) and NOS inhibitor N omega-Nitro-L-arginine methyl ester (L-NAME; ab120136) were from Abcam (Cambridge, UK). Caspase 9 inhibitor Z-LEHD-FMK TFA (S7313) and mitoQ10 (mesylate; S8978) were from Selleckchem (Houston, TX, USA).

### Cell culture

Myoblast lines were cultured in Skeletal Muscle Cell Growth Medium (Promocell, Heidelberg, Germany) supplemented with 20% foetal bovine serum (FBS; ThermoFisher Scientific, MA, USA), 50 μg/mL bovine fetuin, 10 ng/mL recombinant human epidermal growth factor, 1 ng/mL recombinant human basic fibroblast growth factor, 10 μg/mL human recombinant insulin, 0.4 μg/mL dexamethasone (added as PromoCell SupplementMix) and 50 μg/mL gentamycin (Sigma-Aldrich) in a humidified incubator at 37^◦^C with 5% CO_2_. Myoblast were kept sub-confluent, and passaged at ∼60-70% confluency. To induce differentiation, myoblasts were washed twice with phosphate buffered saline (PBS) and cultured in Dulbecco’s Modified Eagle Medium (DMEM) GlutaMax (ThermoFisher Scientific) with 0.5% FBS, 10 μg/mL recombinant human insulin (Sigma-Aldrich) and 50 μg/mL gentamycin for 3-4 days.

Immortalised FSHD patient-derived cellular models were: (i) isogenic ‘54’ derived from the biceps of a mosaic FSHD1 patient [65], with uncontracted control clone 54-6 (13 D4Z4 repeats) and FSHD clone 54-12 (3 D4Z4 repeats); (ii) isogenic ‘K’ (KM271FSH44TA) from the tibialis anterior of a mosaic FSHD1 patient, with uncontracted control clone 44-4 (K4) and FSHD clone 44-8 (K8); and (iii) sibling-matched immortalised model derived from biceps muscle [66], with D4Z4-contracted FSHD clone 16A and uncontracted control clone 16U.

The inducible DUX4 myoblast line LHCN-M2-iDUX4 (‘iDUX4’) was generated on the human LHCN-M2 myoblast line background, where *DUX4* is induced by DOX [38]. The inducible iDUX4^ZSCAN4-tdT^ reporter line was generated by transduction of iDUX4 myoblasts with a Lentiviral vector encoding tdTomato with nuclear localisation signals on both ends under control of the minimal promoter of ZSCAN4 (Fig. S6). pGL3 Zscan4 promoter plasmid was a gift from Stephen Tapscott (Addgene plasmid #69249; http://n2t.net/addgene:69249; RRID: Addgene_69249) [67]. Lentiviral vector was designed with and purchased from VectorBuilder Inc. (Chicago, IL, USA), and lentiviral particles were produced in HEK293FT cells (R70007; ThermoFisher Scientific) as described [68]. Following transduction, myoblasts were selected with Blasticidine S (5 μg/mL) for 2 weeks.

For generation of ρ^0^-like iDUX4 myoblasts (‘iDUX4-ρ^0^’) we used a modified EtBr-based protocol [69]. Briefly, iDUX4 myoblasts were cultured with 50, 100 or 150 nM EtBr with Uridine (100 mg/mL) for two weeks, yielding iDUX4-ρ^0^ myoblasts with varying degrees of mitochondrial impairment. After withdrawal of EtBr treatment, iDUX4-ρ^0^ myoblasts were cultured in full Skeletal Muscle Cell Growth Medium supplemented with 100 mg/mL Uridine.

### Seahorse respirometry

Oxygen consumption rate (OCR), a readout for mitochondrial respiratory chain function, and extracellular acidification rate (ECAR), a measure of lactic acid production, were quantified on a Seahorse XF Pro Analyzer (Agilent Technologies, Santa Clara, CA, USA). For OCR/ECAR measurements, 20000 myoblasts were seeded into a Seahorse XF Cell Culture 96-well plate (102416-100, Agilent Technologies), and respirometry performed 24h later, or differentiation induced for 3-4 days and subsequently analysed. Measurements were performed in Seahorse XF DMEM pH 7.4 Assay Medium (103575-100, Agilent Technologies), supplemented with 10mM Glucose, 1mM pyruvate and 2mM glutamine.

To assay mitochondrial function, cell culture medium was replaced with Seahorse Assay Medium followed by incubation at 37°C for 30 min in a non-CO_2_ incubator. OCR (pmol (O_2_)/min) and ECAR (pH/min) were measured simultaneously at baseline, and after sequential injection of inhibitors/uncouplers. Each measurement cycle consisted of three longitudinal OCR/ECAR measurements for 3 min each, with a 3 min mixing interval between each measurement. After OCR/ECAR measurements at baseline, Oligomycin (1.5μM final concentration), BAM15 (1μM) and Rotenone/Antimycin A (0.5μM each), prepared in Seahorse XF DMEM pH 7.4 Assay Medium, where injected sequentially, and OCR/ECAR recorded after each injection. Seven mitochondrial bioenergetic parameters were calculated from the respiratory profile: basal respiration (respiration at baseline minus non-mitochondrial O_2_ consumption), maximal respiration (maximum uncoupled respiration minus non-mitochondrial O_2_ consumption), ATP-linked respiration (basal respiration minus proton leak), proton leak (Oligomycin-inhibited respiration minus non-mitochondrial O_2_ consumption), coupling efficiency (ratio between ATP-linked respiration and basal respiration in %), spare respiratory capacity (maximal respiration minus basal respiration) and non-mitochondrial respiration (minimum OCR measurement after full inhibition of the respiratory chain).

For quantitation of ATP production rates, OCR/ECAR was monitored at baseline, and after sequential injection of Oligomycin (1.5μm final concentration) and Rotenone/Antimycin A (0.5μM each). Rate of ATP production associated with mitochondrial OXPHOS (mitoATP; pmol O_2_/min) was calculated from OCR with the following formula: (OCR measurement before oligomycin injection minus minimum OCR measurement after Oligomycin injection) x 2 x P/O, where a P/O value (moles of ADP phosphorylated to ATP per atom of Oxygen) of 2.75 was used, as per manufacturer’s instructions. Rate of ATP production associated with conversion of glucose to lactate (glycoATP; pmol ATP/min) is equivalent to glycolytic proton efflux rate (glycoPER; pmol H^+^/min), calculated from ECAR. Total ATP production rate was calculated as the sum of mitoATP and glycoATP production rates [70].

For normalisation of OCR/ECAR to cellular input 5 μg/mL HOECHST33342 (H3570, ThermoFisher Scientific) was added to the last injection, and input cell numbers quantified as fluorescent nuclei counts on a BioTek Cytation 5 Cell Imaging Multimode Reader (Agilent Technologies) coupled to the Seahorse XF Pro Analyzer. Data quality controls, assay parameter calculations and normalisations were performed with Seahorse XF Wave Pro software (Agilent Technologies).

### ROS and ΔΨm measurements

25000 myoblasts were seeded per well into black, clear-bottom polystyrene 96-well plates (ViewPlate®, 6005225; Revvity Inc., Waltham, MA, USA) and ROS levels and ΔΨm quantified 24h later. For myotubes, 40000 myoblasts were seeded per well, induced to differentiate 24h later, and ROS levels and ΔΨm determined after 3-4 days of differentiation. Fluorescent ROS/ΔΨm probes were applied in serum/supplement-free Skeletal Muscle Cell Growth Medium (Promocell) for myoblasts, or DMEM GlutaMax (ThermoFisher Scientific) for myotubes. Prior to assaying, cells were washed twice with 1x Hank’s Balanced Salt Solution (HBSS; with Ca^2+^ and Mg^2+^; Sigma Aldrich), followed by incubation with the ROS/ΔΨm probe for 25 min in the dark at 37°C, 5% CO_2_. General ROS were measured with CM-H_2_DCFDA (final concentration 2.5 μM; C6827, ThermoFisher Scientific) and DNA quantitation performed by simultaneous incubation with Hoechst 33342 (final concentration 0.5 μg/mL) for normalisation to cell input. mitoROS were measured with MitoTracker Red CM-H_2_XRos (final concentration 2.5 μM; M7513, ThermoFisher Scientific) and simultaneous quantitation of mitochondrial content with MitoTracker Deep Red FM (final concentration 250 nM; M22426, ThermoFisher Scientific) and DNA quantitation with Hoechst 33342 (0.5 μg/mL) performed for normalisation. ΔΨm was measured with tetramethylrhodamine methyl ester (TMRM; final concentration 100 nM; T668, thermoFisher Scientific) with simultaneous quantitation of mitochondrial content/DNA for normalisation. After incubation with respective ROS/ΔΨm probes, cells were washed with 1x HBSS, and fluorescence intensities measured on a ClarioStar microplate reader (BMG Labtech, Ortenberg, Germany) in spectral well averaging scan mode (100 measurements per well, scan diameter 6 mm). mitoROS and ΔΨm probe fluorescence was normalised to mitochondrial content (MitoTracker Deep Red fluorescence) after normalisation to input cell quantity (HOECHST33342 fluorescence), as simultaneously assessed via the respective fluorescence intensities in the same well. ROS probe fluorescence was normalised to input cell quantity (HOECHST33342 fluorescence). (mito)ROS and ΔΨm fluorescence measurements are presented as fold change of relative fluorescence intensities after normalisation between controls and samples.

### Apoptosis and caspase activity assays

Apoptosis was measured by the non-invasive, luminescent RealTime-Glo^TM^ Annexin V Assay (JA1001; Promega, Madison, WI, USA). Briefly, 10000 myoblasts were seeded into white, clear-bottom polystyrene 96-well plates (ViewPlate; 6005181, Revvity Inc.). 24h later, assay substrates were added sequentially, according to the manufacturer’s instructions, and the bottom of the plate covered with a white adhesive seal (BackSeal; 6005199, Revvity Inc.) to enhance luminescence signal intensity. A baseline measurement was taken and luminescence signal intensity monitored for up to 48h. For measurements in iDUX4 cells, assay substrates were added with DOX administration. Longitudinal apoptosis measurements are presented as fold change of Annexin V luminescence signal intensity to the baseline measurement.

Caspase 3/7 and 9 activities were measured with the luminescent Caspase-Glo® 3/7 (G8091, Promega) and Caspase-Glo® 9 (G8211, Promega) assays. 25000 myoblasts were seeded into white, clear-bottom polystyrene 96-well plates (Revvity Inc.) and caspase activity measured 24h later. For myotubes, 40000 myoblasts were seeded, induced to differentiate 24h later, and caspase activity measured after 3-4 days. To determine caspase activity, cell medium was removed, the plate covered with a white adhesive seal (Revvity Inc.) and cells lysed with 50 μL of Caspase-Glo® Reagent, prepared freshly according to manufacturer’s instructions. Optimised incubation for maximum signal were 15 min for Caspase 3-7 activity, and 5 min for Caspase 9 (room temperature/shaker/dark). Annexin V and Caspase activity luminescence intensity measurements were performed on a Mithras LB940 multimode microplate reader (Berthold Technologies, Bad Wildbad, Germany) at 0.4s exposure.

### Immunofluorescence microscopy

For immunolabelling, cells were fixed with 4% paraformaldehyde/PBS (J61899-AP, ThermoFisher Scientific) for 15 min, washed 3×5 min in PBS, and permeabilised with 0.1% Triton-X-100/PBS (HFH10, Sigma Aldrich) for 15 min. After 3×5 min washes in PBS, blocking was performed in 5% normal goat serum (GS)/PBS (10000C, ThermoFisher Scientific) for 60 min. After 3×5min washes with PBS, primary antibody was added in 1% GS/PBS on a rocker overnight at 4°C. Primary antibodies were: mouse monoclonal anti-MyHC (1:400; MF-20, Developmental Studies Hybridoma Bank, Iowa City, IA, USA), mouse monoclonal anti-tubulin beta (1:400; E7, DSHB), mouse monoclonal anti-DUX4 (1:500; clone 9A12; MABD116, Merck Millipore, Croxley Park, UK), rabbit monoclonal anti-DUX4 (1:1000; E5-5; ab124699, Abcam) and rabbit polyclonal anti-tdTomato (1:500; TA150128, ThermoFisher Scientific). Cells were then washed 3×5 mins in PBS, and incubated with secondary antibody in 1% GS/PBS for 60 min in the dark at room temperature. Secondary antibodies were: goat anti-mouse IgG (H + L) AlexaFluor-488 (1:500, A-11001, ThermoFisher Scientific) and goat anti-rabbit IgG (H + L) AlexaFluor-594 (1:500, A-11012, ThermoFisher Scientific). After 3×5 min washes with PBS, nuclei were stained with 0.5 μg/mL HOECHST33342 in PBS for 10 min, samples washed again with PBS and imaged using Invitrogen EVOS™ M5000 Imaging (ThermoFisher Scientific).

### Reverse transcription and quantitative PCR

Total RNA was extracted with the RNeasy Mini Kit (74104, QIAGEN, Venlo, Netherlands) and quantified on a NanoDrop ND-1000 UV/Vis spectrophotometer (ThermoFisher Scientific). 1 μg of RNA was used for cDNA synthesis using the QuantiTect Reverse Transcription Kit (205313, QIAGEN). RT-qPCR analyses were in triplicate from 5 ng input cDNA on a ViiA7 thermal cycler (Applied Biosystems, Warrington, UK), using Takyon® Low ROX SYBR 2X MasterMix blue dTTP (UF-LPMT-B0701, Eurogentec, Seraing, Belgium) as per manufacturer’s instructions. Target gene expression was normalised to the housekeeper *RPLP0*, and values represented as relative expression (2^-ΔCT^). Primer sequences (from Sigma Aldrich): *RPLP0*-fwd: 5’-TGGTCATCCAGCAGGTGTTCGA-3’, *RPLP0*-rev: 5’-ACAGACACTGGCAACATTGCGG-3’, *ZSCAN4*-fwd: 5’-TGCCTCCTGGATTCAAACA-3’, *ZSCAN4*-rev: 5’-AGTGTTCTATACCATCACTGGTCCT-3’, *PRAMEF1*-fwd: 5’-GCTGCTTACACAGAAGGACCG-3’, *PRAMEF1*-rev: 5’-CTCTGGGAAGCAGGACAGG-3’, *TRIM43*-fwd: 5’-CAGGAACTAATGGAAATGTGTCAT-3’, *TRIM43*-rev: 5’-GACTCACTCCTTGCCACGAT-3’.

### Quantification of relative mitochondrial DNA content

Total cellular DNA was extracted with the QIAamp DNA Mini Kit (QIAGEN) as per manufacturer’s instructions and quantified on a NanoDrop ND-1000 UV/Vis spectrophotometer (ThermoFisher Scientific). Relative amounts of mitochondrial (housekeeper *tRNA^Leu(UUR)^*) and nuclear (housekeeper *β-2-microglobulin; β2M*) DNA were determined in quadruplicate with qPCR on a ViiA7 thermal cycler (Applied Biosystems) from 5 ng input DNA using Takyon® Low ROX SYBR 2X MasterMix blue dTTP (Eurogentec). Primer sequences (from Sigma Aldrich): *β2M*-fwd: 5’-TGCTGTCTCCATGTTTGATGTATCT-3’, *β2M*-rev: 5’-TCTCTGCTCCCCACCTCTAAGT-3’, *tRNA^Leu(UUR)^*-fwd: 5’-CACCCAAGAACAGGGTTTGT-3’, *tRNA^Leu(UUR)^*-rev: 5’-TGGCCATGGGTATGTTGTTA-3’ [71]. Relative mtDNA content was calculated by 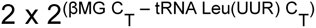 [72], and presented as fold change after normalisation between controls and samples.

### Transcriptomic analyses

Raw RNA-seq read counts and matched metadata information were acquired from the Gene Expression Omnibus (accession number GSE140261 [47]). Disease severity grouping (lowest disease severity: G1 to highest disease severity: G4) was extracted for contrasting with no factoring of age or gender, as no gender/age data was available for controls. For differential expression analysis, un-normalised gene-level counts were imported to DESeq2 [73] and genes with a total count <10 across all samples were filtered out. A DESeq2 object was constructed based on the design ∼ disease severity grouping (Control, G1, G2, G3, G4) with pairwise contrasts of Control vs G1-G4 disease severity groups. Gene annotation used transcript ID annotation through the org.Hs.eg.db package in Bioconductor to acquire gene names.

Enrichment and over-representation analysis (ORA) was performed using MSigDB HALLMARK [74] and Gene Ontology:Biological Processes (GO:BP; Gene Ontology Consortium) gene sets, retrieved via msigdbr (CRAN package). ORA as universal enrichment was performed on upregulated (log2FC>0.5) and downregulated (log2FC<-0.5) differentially expressed genes (DEGs) using the package clusterProfiler::enricher [75] with all gene ontology HALLMARK and GO:BP gene sets investigated. Multiple testing was performed using Benjamini–Hochberg false discovery rate (FDR) and was controlled across tested gene sets within each collection and contrast.

Plotting was performed using ggplot2 (https://ggplot2.tidyverse.org). Analyses were performed in R (version 4.5.1; 2025-06-13) using Rstudio (2025.5.1.513) with the following packages: readr (version 2.1.5), dplyr (version 1.1.4), tidyr (version 1.3.1), ggplot2 (version 3.5.2), ggpubr (version 0.6.0), ggrepel (version 0.9.6), DESeq2 (version 1.48.1), clusterProfiler (version 4.16.0), org.Hs.eg.db (version 3.21.0) and msigdbr (version 25.1.1).

### Statistical analysis

Statistical analysis was performed using GraphPad Prism version 10.5.0 for Windows (GraphPad Software, San Diego, CA, USA; www.graphpad.com). Heatmap for Apoptosis dose-response matrices and HAS synergy scores were made with the SynergyFinder+ web application [76].

Experiments were performed at least 3 independent times, with N numbers and technical replicates given in each figure. Variance between groups was compared using a Brown Forsythe test and revealed no significant difference. Comparison between two groups was using an unpaired homoscedastic two-tailed student’s *t*-test. Comparison of more than two groups was using a one-way analysis of variance (ANOVA) followed by either Dunnett’s post-test when different groups were compared with the control group or Tukey’s post-test when different groups were compared with each other. *p <* 0.05 was considered significantly different, with *p* values for significant comparisons in figures.

## Supporting information

Heher et al_Supplementary Information

## DATA AVAILABILITY

GSE140261 is publicly available. Other raw and source data can be accessed on reasonable request.

## ACKNOWLEDGEMENTS

We thank Michael Kyba (University of Minnesota, Minneapolis, MN, USA) for the LHCN-M2-iDUX4 human myoblast line, Vincent Mouly (Center for Research In Myology, UMRS 974 Sorbonne Université-INSERM, Paris, France) and Charles P. Emerson (Wellstone Muscular Dystrophy Program, University of Massachusetts Medical School, Worcester, MA, USA) for immortalised myoblast lines. We are grateful to Alexandra Belayew, Andrey Kozlov, Simon Hughes and Dustin Bagley for helpful discussion, and to Nick Howe for assistance with Seahorse respirometry.

## CONTRIBUTIONS

PH, CRSB, JG and PSZ secured funding, conceptualised research, interpreted data and wrote the manuscript. PH and MG generated cell models and designed experimental approaches, and, together with JPV, LG, MT and JP, conducted experiments, acquired and analysed data. RF, ENE and CRSB processed RNA-sequencing data and performed transcriptomic analyses on biopsies with support from GT, who secured funding for RF and currently for PH.

