## Supplementary material for "Mitochondrial Respiratory Chain Function is crucial for Muscle Toxicity in Facioscapulohumeral Muscular Dystrophy": Heher et al_Supplementary Information

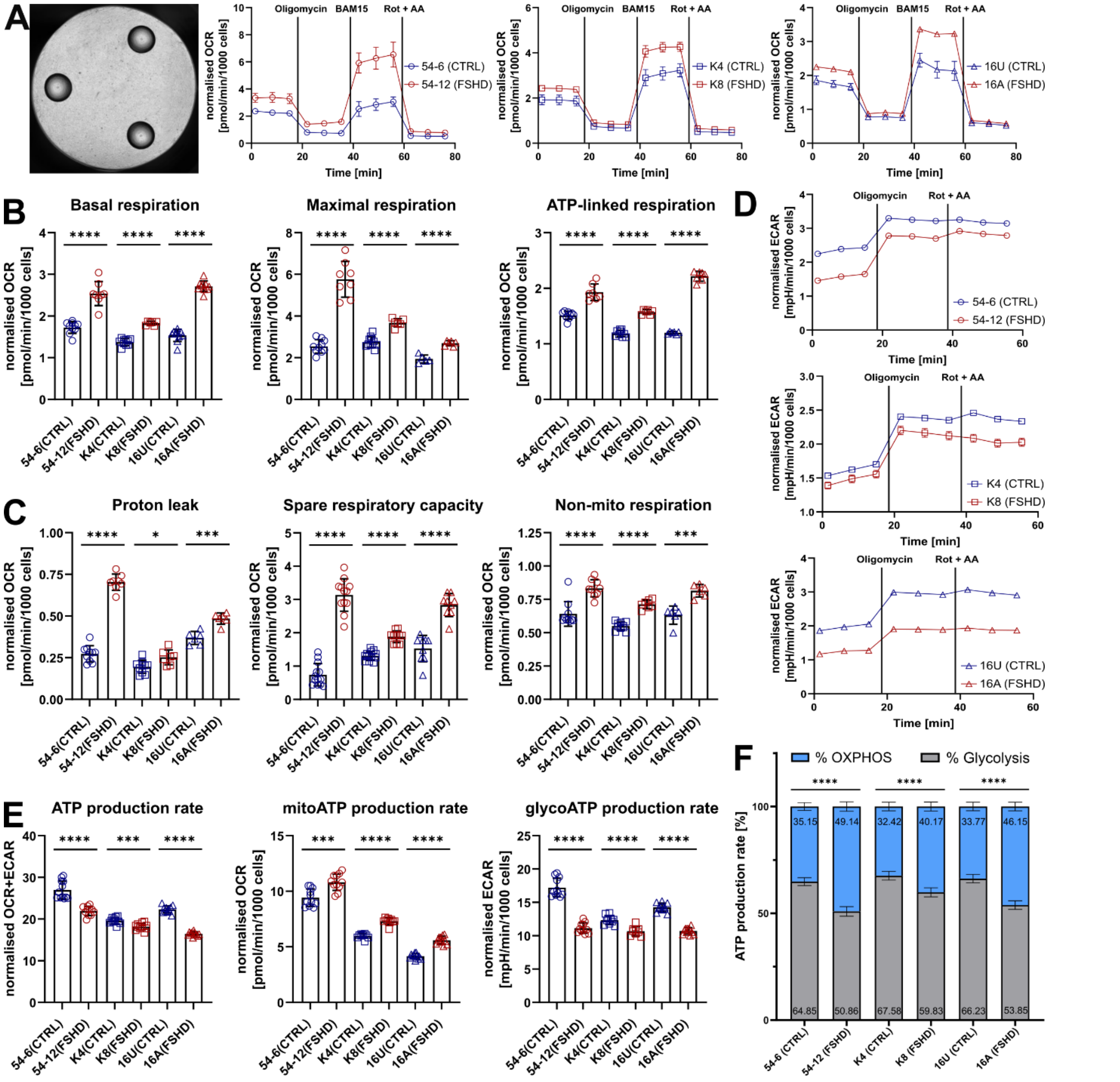


**Figure S1:** ***FSHD myoblasts are characterised by an ATP deficit and altered metabolic setup*** **(A)** Left: Brightfield micrograph of an FSHD patient-derived myoblast culture after Seahorse respirometry. Right: normalised oxygen consumption rate (OCR) curves of three independent FSHD myoblast cell models (54-12, K8, 16A), alongside matched unaffected controls (54-6, K4, 16U). **(B)** Increased mitochondrial basal, maximal and ATP-linked respiratory function in FSHD myoblasts. **(C)** Increased proton leak, spare respiratory capacity and non-mitochondrial ("Non-mito") respiration in FSHD myoblasts. **(D)** Normalised extracellular acidification rate (ECAR) curves of three FSHD myoblast cell models, alongside matched unaffected controls. **(E)** ATP production rates demonstrating an ATP deficit in FSHD myoblasts arising from defective glycolysis (glycoATP), resulting in **(F)** an altered balance between mito and glycoATP production. (n=6-10, data is mean ± s.d., where **p*<0.05, ****p*<0.001, *****p*<0.0001).

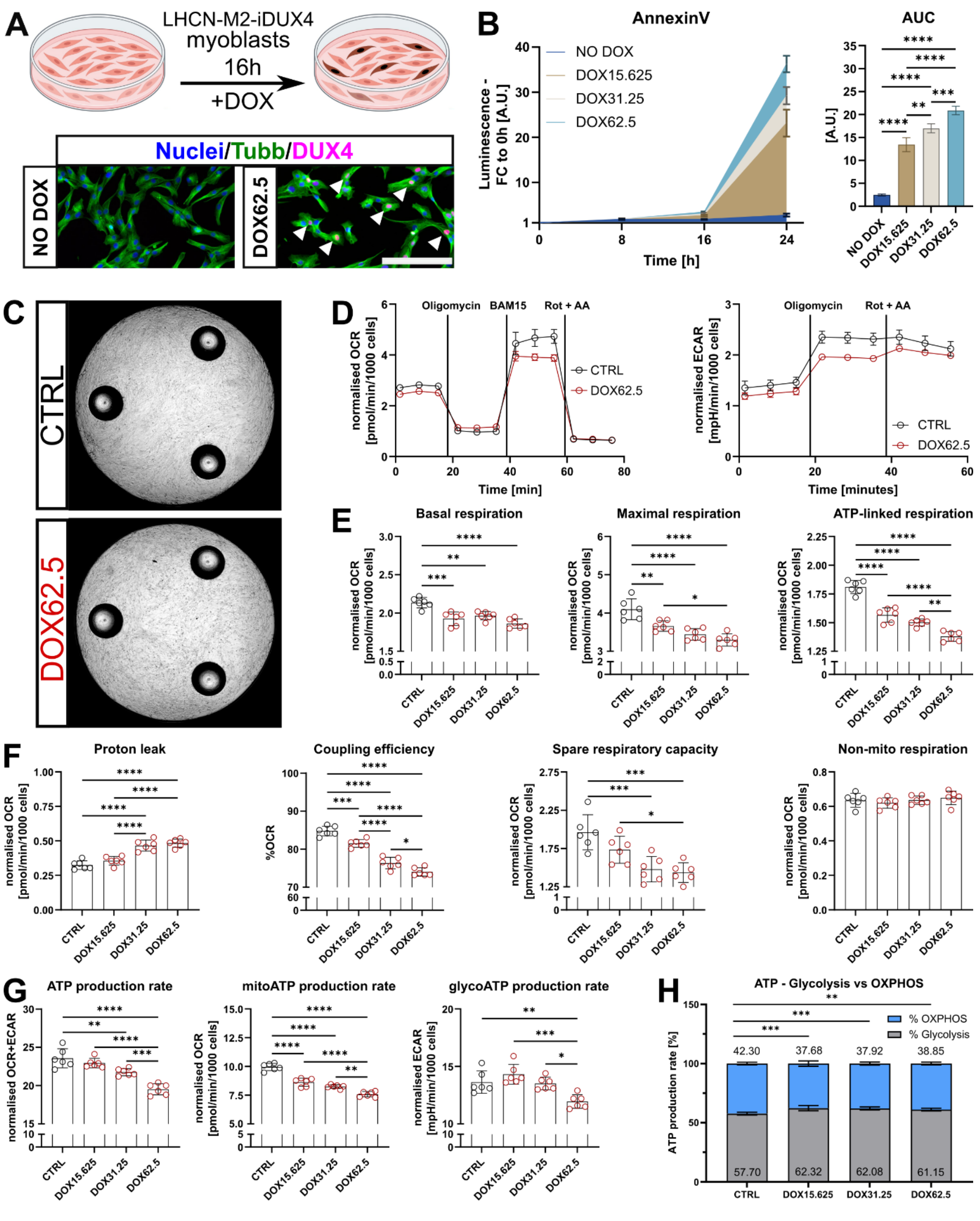


**Figure S2: *DUX4 expression in human myoblasts triggers metabolic stress-induced apoptosis through mitochondrial dysfunction in a dose-dependent manner***

**(A)** Top: schematic of the LHCN-M2-DUX4-inducible human myoblast model (iDUX4), where *DUX4* expression can be induced through doxycycline (DOX). Scheme was created with BioRender.com. Bottom: immunofluorescence micrographs of iDUX4 myoblasts expressing DUX4 after DOX (62.5 ng/mL) administration for 16h (pink=DUX4, green=β-Tubulin, blue=nuclei; scale bar=200 μm). Arrowheads mark DUX4-positive nuclei. **(B)** Dose-dependent increase of apoptosis in response to varying levels of DUX4 induction for 24h, as assessed by longitudinal Annexin V luminescence assaying (left) with subsequent area under curve (AUC) analysis (right). **(C)** Brightfield micrographs of iDUX4 control (CTRL, top) and DOX-induced (62.5 ng/mL for 16h, bottom) myoblasts after Seahorse respirometry. **(D)** Normalised oxygen consumption rate (OCR - left) and extracellular acidification rate (ECAR - right) curves of iDUX4 control (CTRL) and DOX-induced (62.5 ng/mL DOX for 16h) myoblasts. **(E)** Dose-dependent reduction of mitochondrial basal, maximal and ATP-linked respiratory function in iDUX4 myoblasts after induction of DUX4 expression through exposure to different concentrations of DOX for 16h. **(F)** Increased proton leak and reduced coupling efficiency and spare respiratory capacity, but unchanged non-mitochondrial ("Non-mito") respiration in iDUX4 myoblasts after exposure to varying levels of DUX4 induction through DOX for 16h. **(G)** ATP production rates demonstrating a gradual ATP deficit in iDUX4 myoblasts after exposure to varying levels of DUX4 induction through DOX for 16h, arising mainly from defective oxidative phosphorylation (OXPHOS; mitoATP), resulting in **(H)** an altered balance between mito and glycoATP production. (n=5-6, data is mean ± s.d., where *$p$<0.05, **$p$<0.01, ***$p$<0.001, ****$p$<0.0001).

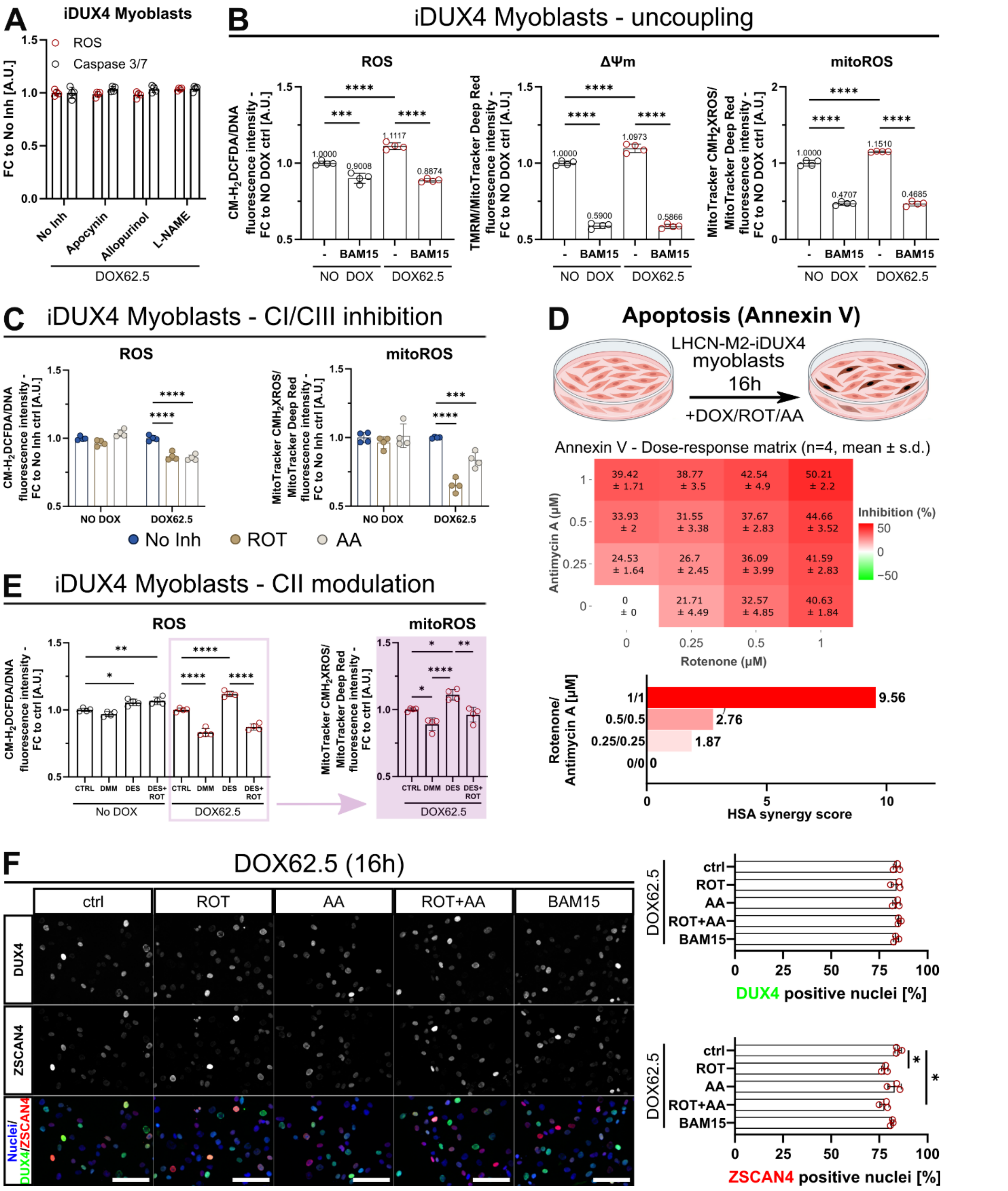


**Figure S3:** ***DUX4 causes oxidative stress-induced apoptosis via Caspase 9 through reverse electron transfer (RET)-ROS release from the respiratory chain in dysfunctional mitochondria***
**(A)** ROS levels and Caspase 3/7 activation in iDUX4 myoblasts after DUX4 induction with DOX (62.5 ng/mL) for 16h are not reduced when NOX (Apocynin 10μM), XO (Allopurinol 10μM) or NOS (L-NAME 5mM) are inhibited with DOX for 24h. **(B)** ROS levels, ΔΨm and mitoROS levels are higher in iDUX4 myoblasts after DUX4 induction with DOX (62.5 ng/mL) for 16h, with a greater reduction in DUX4-expressing myoblasts compared to controls (NO DOX) after uncoupling of the mitochondrial respiratory chain with BAM15 (1 μM for 25 min). **(C)** ROS and mitoROS levels in iDUX4 myoblasts after DUX4 induction with DOX (62.5 ng/mL) for 16h are uniquely reduced in DUX4-expressing

myoblasts after complex I (CI) inhibition with ROT (0.5 µM for 25min) or complex III (CIII) inhibition with AA (0.5 µM for 25 min). **(D)** Top: Apoptosis (Annexin V) inhibition dose-response matrix of iDUX4 myoblasts after induction of DUX4 with DOX (62.5 ng/mL) for 16h with simultaneous ROT (0.25, 0.5, 1 µM) versus AA (0.25, 0.5, 1 µM) supplementation - heatmap of apoptosis inhibition revealing a synergistic effect of combinational administration of ROT and AA. Scheme was created with BioRender.com. Bottom: Highest Single Agent (HAS) synergy scores indicating an increase in synergy of ROT and AA with increasing concentrations. **(E)** ROS levels in iDUX4 myoblasts after DUX4 induction with DOX (62.5 ng/mL) for 16h with subsequent modulation of complex II (CII)-linked respiration through administration of DMM (10 mM for 25 min), DES (5 mM for 25 min), or DES (5 mM for 25 min) and ROT (0.5 µM for 25 min). Modulation of CII-linked respiration after DUX4 expression affects ROS levels by gauging mitoROS levels from the respiratory chain (purple insert). **(F)** Immunofluorescence micrographs of iDUX4$^{ZSCAN4\text{-}tdT}$ reporter myoblasts 16h after exposure to DOX (62.5 ng/mL) with concomitant administration of ROT (0.5 µM), AA (0.5 µM), ROT+AA (0.5 µM each) or BAM15 (1 µM) shows no reduction in DUX4$^+$ nuclei, but moderate reduction of ZScan4$^+$ nuclei when CI is inhibited with ROT alone, or in combination with AA [green=DUX4, red=tdTomato (ZSCAN4), blue=nuclei; scale bar=100µm]. (n=3-4, data is mean ± s.d., where *$p$<0.05, **$p$<0.01, ***$p$<0.001, ****$p$<0.0001).

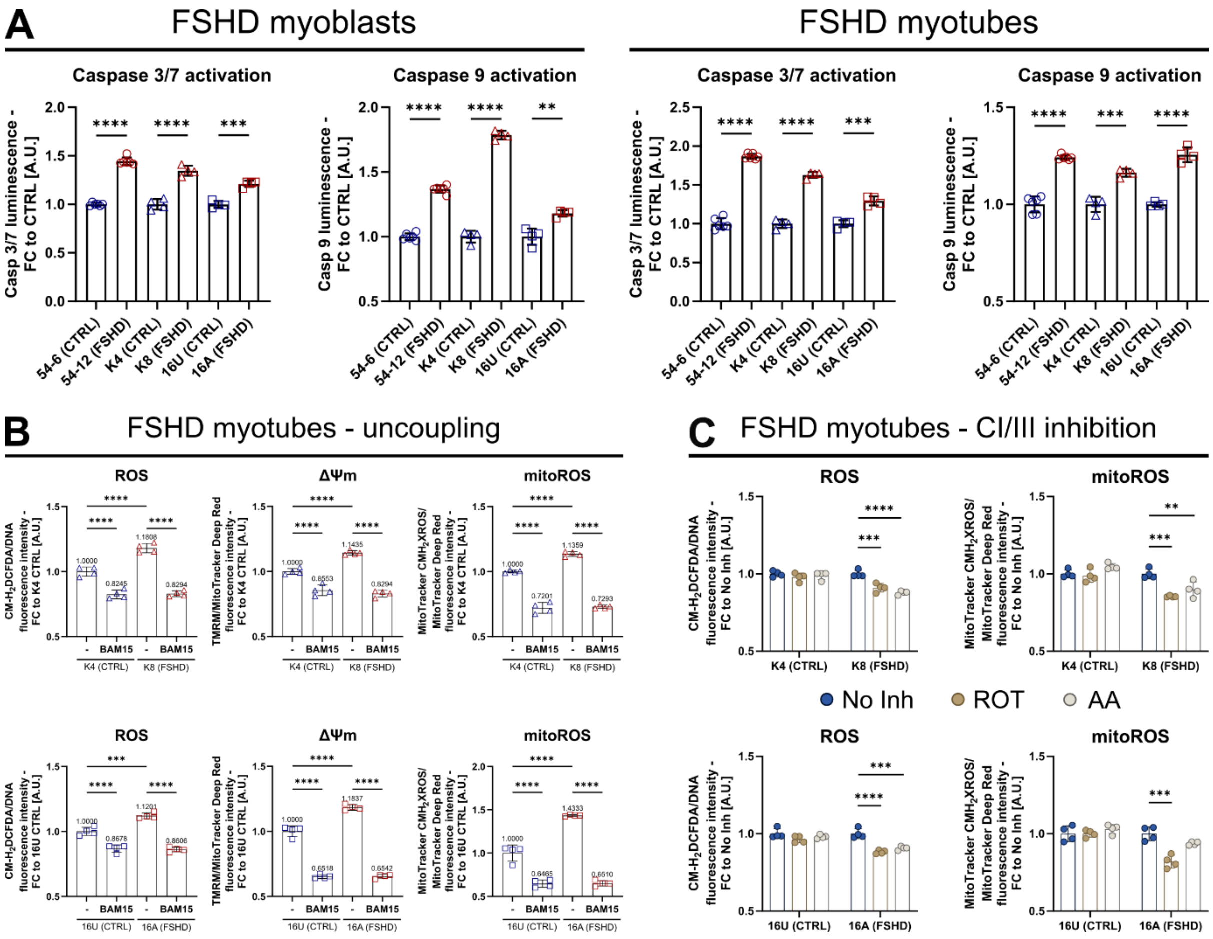


**Figure S4:** ***Dysfunctional mitochondria are the main ROS producing system in FSHD muscle cells and trigger apoptosis in a Caspase 9-dependent manner***

**(A)** Higher baseline Caspase 3/7 and Caspase 9 activity in 54-12, K8, 16A FSHD myoblasts (left) and myotubes (right), compared to unaffected 54-6, K4, 16U controls. **(B)** ROS levels, ΔΨm and mitoROS levels are higher FSHD myotubes (K8: top; 16A: bottom), with a greater reduction in FSHD myotubes compared to controls (K4 and 16A, respectively) after uncoupling of the mitochondrial respiratory chain with BAM15 (1 µM for 25 min). **(C)** ROS and mitoROS levels are reduced in FSHD myotubes (K8: top; 16A: bottom) after complex I (CI) inhibition with ROT (0.5 µM for 25 min) or complex III (CIII) inhibition with AA (0.5 µM for 25 min), but not in controls (K4 and 16A, respectively). (n=4, data is mean ± s.d., where ****p*<0.01, *****p*<0.001, ******p*<0.0001).

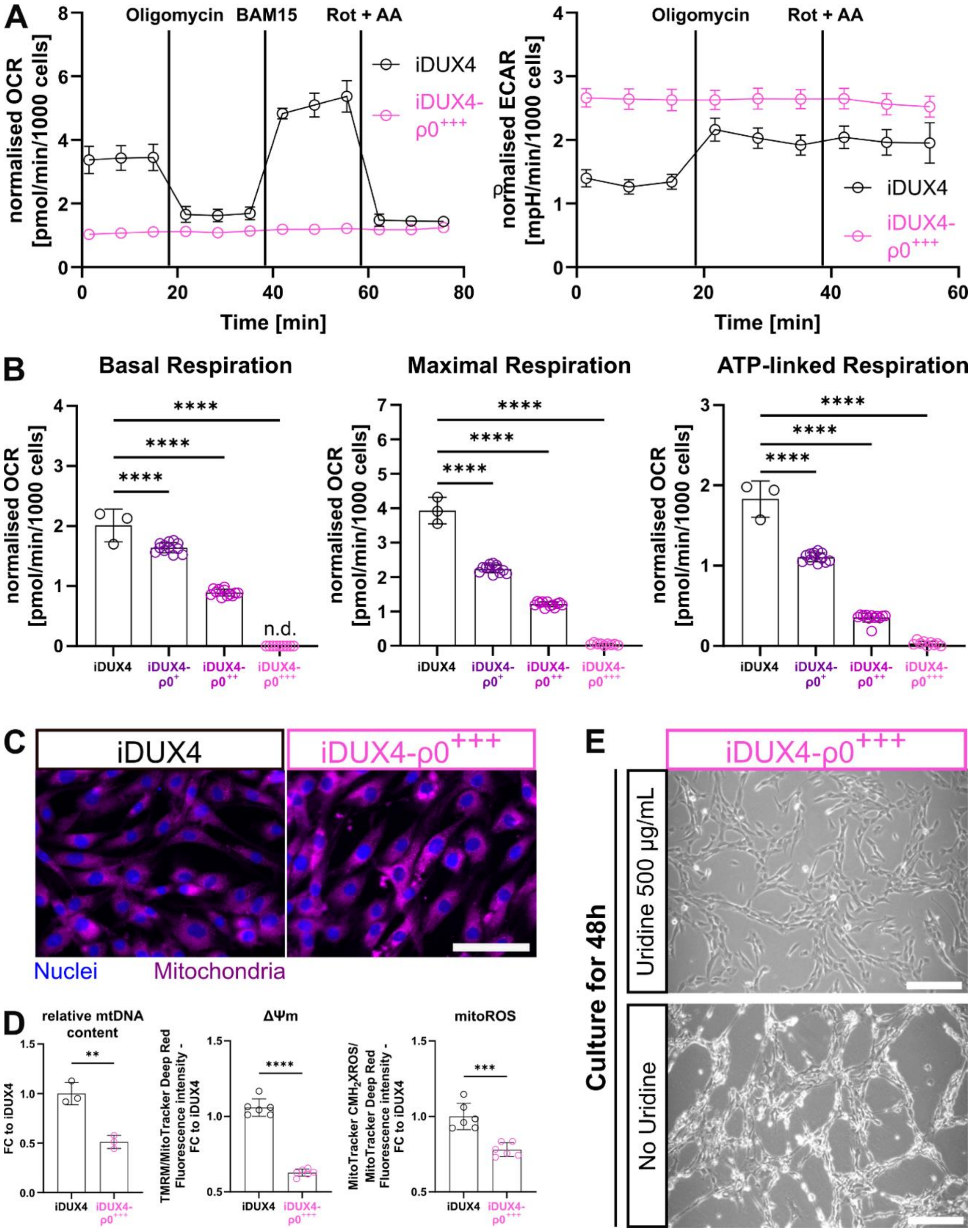


**Figure S5:** ***Characterisation of iDUX4- $\rho^0$ myoblast mitochondria***

**(A)** Normalised oxygen consumption rate (OCR - left) and extracellular acidification rate (ECAR - right) curves of unmodified iDUX4 versus iDUX4-$\rho^{0+++}$ myoblasts. **(B)** Increased impairment of basal, maximal and ATP-linked mitochondrial respiratory function in iDUX4- $\rho^{0+}$ and iDUX4-$\rho^{0++}$ and full OXPHOS inhibition in iDUX4-$\rho^{0+++}$ myoblasts compared to unmodified iDUX4 controls. **(C)** Representative fluorescence micrographs [purple=MitoTracker Deep Red FM (mitochondria), blue=nuclei; scale bar=100µm] of unmodified iDUX4 and iDUX4-$\rho^{0+++}$ myoblasts showing no overt difference in mitochondrial mass, shape and distribution. **(D)** Reduced relative mtDNA content, ΔΨm and mitoROS levels in iDUX4-$\rho^{0+++}$ myoblasts compared to unmodified iDUX4 controls. **(E)** Brightfield micrographs of iDUX4-$\rho^{0+++}$ myoblasts cultured with or without Uridine (500 µg/mL) supplementation for 48h (bar=250µm). (n=3-12, data is mean ± s.d., where ***p*<0.01, ****p*<0.001, *****p*<0.0001).

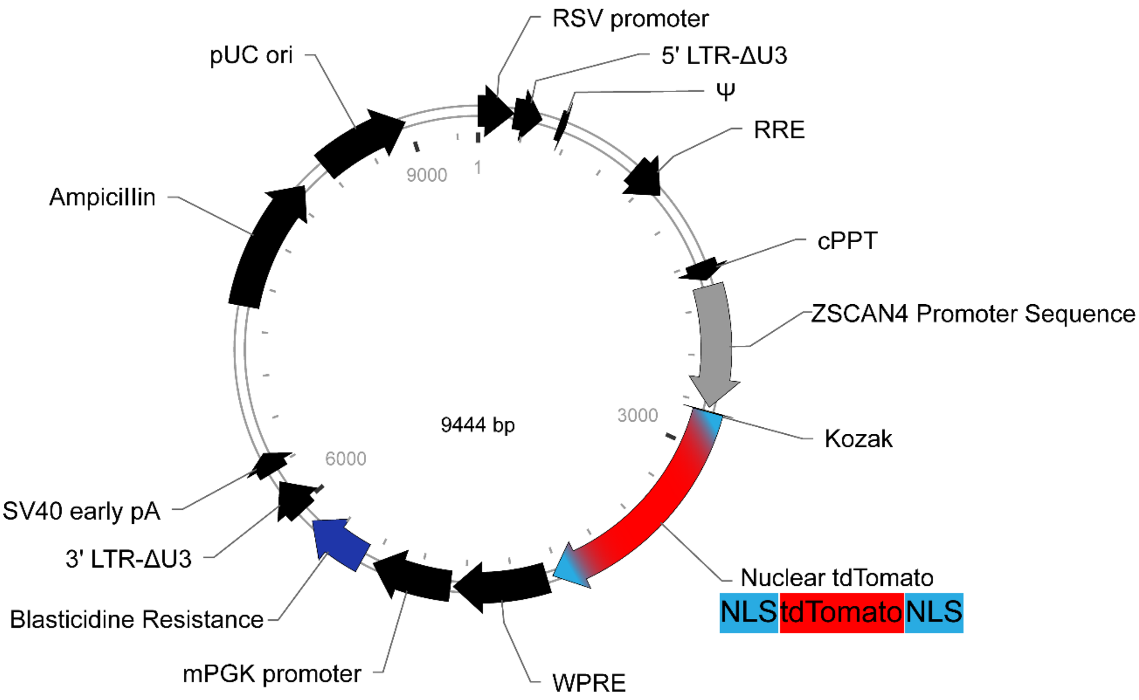


**ZSCAN4 Promoter Sequence:**

CGGTTCAAGTAATTCTCCTGCCTCAGCCTCCCAAGTAGCTGGGATTACAGGTGCCCACCAGCATGCCCGGCTAATTTTTGTA
TTTTTAGTAGAGAAAGGGTTTCACCATGTTGGCCAGGCTGGTCTTGAACTCCTGATCTCAGGTGATCTGCCTGCCTCAGCCT
CCCAAAGTGCTGAGATTACAGGTGTGAGCCACTGCACCTAGCCTGCAGAAATATTAATGTGTTTGTTTATGACCTGTTGACCC
AGGACCAGTGATGGTATAGAACACTGTATTAGAGACATGGAGCTGGGGCTGGATGAAGATTCCATCAGTAATTCAATCAACAG
ACAAGTGTTATCCAATCACGTCTTTAAATCAATCACTGACATGGAGCTGGGGCTGGATGAAGATTCCATCGGTAATTCAATCAA
CAGACAAGTGTTATCCAATCACGTCTTTAAATCAATCACTGATCCCAGCCCCTATAAAAGGGAGCAGCCTTAGGAGGCACATC
AGATAAACCCAGTGTGGAAAGCTAGTCACACATCAGCTCAGTGTTCGGCCCGGGATTACCCAGTCAACCAAGGAGGTAAGCT
TCCATAATGGAAGGAAAATTTTGTGCCTTTGAGTGTGTATACATTTTGTGTCTTTGAGTGTGTATACATAATTTTGTGTCTTTGAG
TGTGTATACATGTGTATGTAGGCATAGCTATTTCAGGAAAATTCCTAACTACAGAGATATGGGGGTCTCCCAGGTGTAGGGAATG
GTCTTTTGTTACAACAGGAGCTC

Putative DUX4 binding sites (Identified with Jaspar Matrix MA0468.1, Threshold profile score 90%)

**Figure S6:** ***Generation of the iDUX4$^{ZSCAN4\text{-}tdT}$ reporter myoblast line***
**Top:** Schematic of the Lentiviral vector encoding for tdTomato with nuclear localisation signals on both ends under control of the minimal promoter region of ZSCAN4.
**Bottom:** Sequence of the minimal ZSCAN4 promoter region, with putative DUX4 binding sites highlighted in purple.
